# In depth characterisation of the tumour microenvironment reveals HHV-8 dependent immune regulation in HIV-associated and classic Kaposi Sarcoma

**DOI:** 10.64898/2025.12.16.694599

**Authors:** Claudia AM Fulgenzi, Alessia Dalla Pria, Yiran Zhao, Alberto Giovanni Leone, Justin Weir, Ignazio Puccio, Krista S Tuohinto, Päivi M Ojala, Mark Bower, David J. Pinato

## Abstract

**Background:** Kaposi’s sarcoma (KS) is the most common malignancy occurring in people living with HIV (PLWH). The clinical course of KS in PLWH established on combined anti-retroviral therapy (cART) resembles that of classic KS. However, no studies have so far been conducted to compare clinical and immunologic characteristics of these subtypes.

**Objectives:** To compare the clinical and biological characteristics of HIV-associated KS in patients with well-controlled viral infection (N=21) and non-HIV associated KS (N=19).

**Methods:** Clinical data were prospectively collected from patients treated at the National Centre for HIV malignancies at Chelsea & Westminster Hospital. Archival tumour samples were retrieved to perform targeted transcriptomic analysis using the Nanostring nCounter platform. The results were validated by multiplex immune-fluorescence.

**Results:** Median progression free survival (mPFS) was comparable between groups. The tumour microenvironment (TME) of non-HIV associated KS was characterized by upregulation of pathways associated with adaptive and innate immunity and angiogenesis. Transcripts relating to glycolysis and endothelial to mesenchymal transition were upregulated in the TME of HIV-associated cases. Furthermore, the TME of HIV-associated cases was associated with lower infiltration of CD4 activated cell. In both cohorts, we found the expression of HHV-8 genes to positively correlate with activated CD4, CD8, NK and immune checkpoints gene expression.

**Conclusions:** In conclusion, we showed that the TME of KS in the presence of well-controlled HIV infection is characterized by a lower degree of inflammation and more pronounced epithelial to mesenchymal transition. We also identified intra-tumoral HHV-8 gene expression as a driver of TME composition.

## Introduction

Kaposi’s sarcoma (KS) is a mesenchymal tumour of possible lymphovascular origin that recognises Human Herpes Virus 8 (HHV-8) infection as the necessary but not sufficient pathogenic mechanism^4^. Uncontrolled HIV infection represents one of the most common causes orf KS development and progression ^5^ and, despite the drop in its incidence after the introduction of combined-antiretroviral therapy (cART), KS remains the most common malignancy in people living with HIV (PLWH).

The current taxonomic classification of KS recognises classic, iatrogenic, epidemic, endemic and KS in men who have sex with men (MSM) as 5 independent classes based on disease epidemiology. Recent evidence from our group has emphasised that the degree of baseline immune suppression, independently from the cause, is the main factor influencing the clinical course of KS^3^, and reducing the degree of immune suppression leads to KS remission.

For instance, while formally included into the “epidemic” group, HIV-associated KS includes clinical entities occurring in the presence of well-controlled HIV, resembling more closely that of classic KS rather than that of AIDS-associated KS. Equally, iatrogenic KS has often a clinical course that mimics that of AIDS-associated KS rather than other forms of KS ^3^. In the case of KS persistence or recurrence, or when the underlying immune suppression cannot be treated, chemotherapy is effectively used to treat KS ^6^. However, while immunosuppression remains key for KS development and progression, little is known about the biological and immunological characteristics of KS occurring in the absence of clinically significant immune suppression.

The strong pathogenic reliance of KS upon defective adaptive immunity makes the use of immunotherapy particularly appealing in this malignancy.

So far, immune checkpoint inhibitors have shown promising efficacy in both classic and HIV-associated KS^7 8^, with an objective response rate reaching 87% for the combination of ipilimumab (anti-CTLA-4) plus nivolumab (anti-PD1) in classic KS^9^. Similarly, immunomodulatory agents such as pomalidomide have shown anti-cancer activity in both HIV and non-HIV cases^10^. A better understanding of the biological mechanisms leading to KS in the absence of clinically significant immune suppressions remains pivotal to develop new treatment strategies.

In this study, we aimed to investigate the differences between HIV-associated KS occurring in the presence of well-controlled HIV infection and non-HIV associated KS without clinically evident immune suppression, by comparing the clinical characteristics and the composition of the tumour microenvironment (TME) of two matched cohorts.

## Methods

### Patient population

The two study cohorts were selected from a prospectively maintained dataset including all patients treated at the National Centre for HIV malignancies at the Chelsea and Westminster Hospital. The study included 40 consecutive patients with histology proven KS diagnosed from 01/2008 to 02/2015 and available tissue samples. Diagnostic tissue was retrieved from the pathology archive. Patients with non-controlled HIV, defined as viral load > 200 copies/ml and CD4 count < 200 cells/mm^3^, or other causes of immune suppression were excluded. Data collection was approved by the National Centre for HIV malignancies and tissue collection was approved the Imperial College Healthcare Tissue Bank (R180090). Baseline characteristics are reported descriptively as percentages (%) for categorial variables and median with interquartile ranges for continuous ones. Categorical variables were compared using the chi-squared test, and continuous variables using the Kruskal-Wallis test. Progression free survival (PFS) was calculated as the time from last treatment (either local or systemic) commencement to death or evidence of KS progression, whichever came first. Patients not reporting progression at the time of data cut-off were censored at the time of last follow-up. Median PFS was estimated using Kaplan-Meier (KM) method, whereas median follow-up times were estimated with the reverse Kaplan-Meier method.

### Transcriptomic profiling

After quality check for the presence of residual tumour, 3 scrolls by 10 microns were cut from formalin-fixed paraffin embedded tumour blocks. RNA was extracted from each sample using the AllPrep DNA/RNA FFPE Kit by Qiagen as per manufacturer instructions. After RNA quantification using the Nanodrop profiler, we performed gene expression profiling using the NanoString nCounter Analysis System with the PanCancer IO 360 Panel, to which a custom CodeSet of 10 HHV-8 genes was added (**Supplementary Tables 1-2**).

### Gene expression data processing and normalization

Details about gene expression analysis are reported in **Supplementary Method 1**.

### Immunofluorescence

Slides prepared from 12 formalin-fixed paraffin-embedded (FFPE) tissue blocks were obtained and processed for both immunohistochemistry (IHC) and multiplex immunofluorescence (mIF) following standard protocols for clinical trial samples developed by the ICR/RMH Integrated Pathology Unit (IPU). Antibody staining was carried out using the BOND RX 3.0 Fully Automated Slide Stainer (Leica Biosystems).

To explore changes in immune cell populations and viral protein expression, 2 multiplex panels comprising 6 biomarkers each plus DAPI were developed using the PhenoImager HT system (Akoya Biosciences). Details about the methods are reported in **Supplementary Methods 2** and **Supplementary Tables 3-4**.

### Plasma HHV-8-DNA quantification

Plasma HHV8 was recorded at the time of tissue biopsy. Plasma HHV8 samples were analysed using a quantitative nested PCR approach, targeting the open reading frame-26 (minor capsid protein) gene of KSHV.

## Results

### Clinical characteristics of classic and HIV-associated KS cohorts

The whole clinical cohort included 40 patients. Among them, 19 had non-HIV associated KS (classic), and 21 HIV-associated KS. Among those with HIV-associated KS, all of them were stablished on cART by at least 3 months at the time of samples collection, had HIV-RNA < 200 copies/ml and all had CD4 count >200 cells/mm^3^. The baseline characteristics of the whole cohort and the two main subgroups are summarised in **Table 1**. In brief, most of the patients were males (N=37), 24 out of 40 had prior loco-regional therapies to treat KS, with excision being the most common (N=17); 22 received at least one line of systemic therapy. The median time from first KS diagnosis (either clinical or histology proven) and the last histology confirming KS (referred to the tissue used for the translational analysis of this study) was 6.8 months (range 0-21 years). In those requiring systemic therapy, all the samples were collected prior to the last line of treatment start. As reported in **Table 1**, patients with HIV-associated KS had younger age at first diagnosis, higher peripheral blood CD8 count and lower HHV-8 levels. Other baseline characteristics were well-balanced across the 2 cohorts. After a median follow-up of 15.3 months (95%CI:12.0-30.3), median PFS in the whole cohort was 17.3 months (95%CI:8.0-NA, **Supplementary Figure 1A**), which was not different when stratified according to the HIV status (HR=1.3; 95%CI: 0.50-3.48, p=0.6, **Supplementary Figure 1B**).

**Figure 1.**
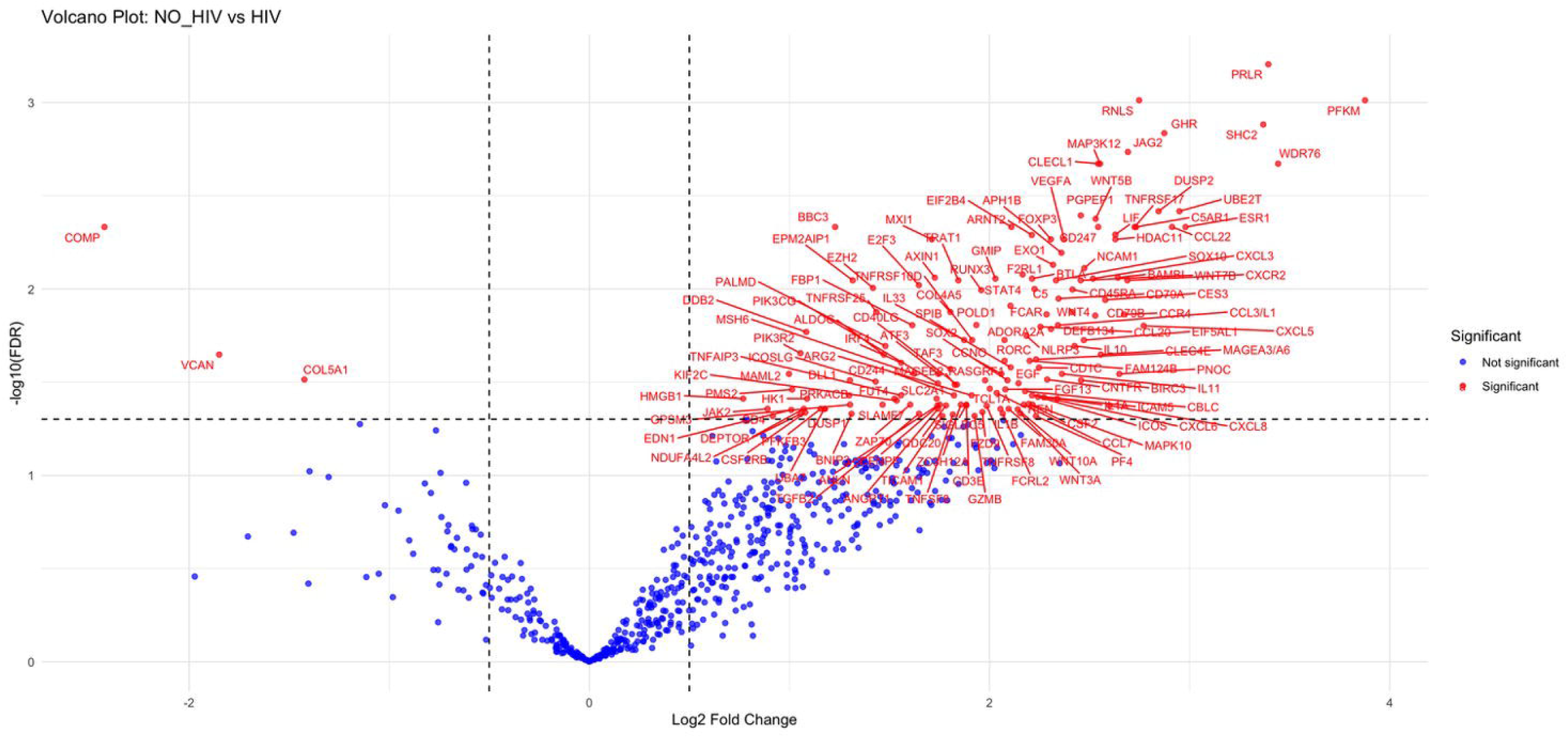
Volcano plot showing the differential gene expression between Non-HIV cases (on the right) and HIV cases (on the left). Red genes represent significantly differentially expressed genes after FDR adjustment.

**Table 1.**
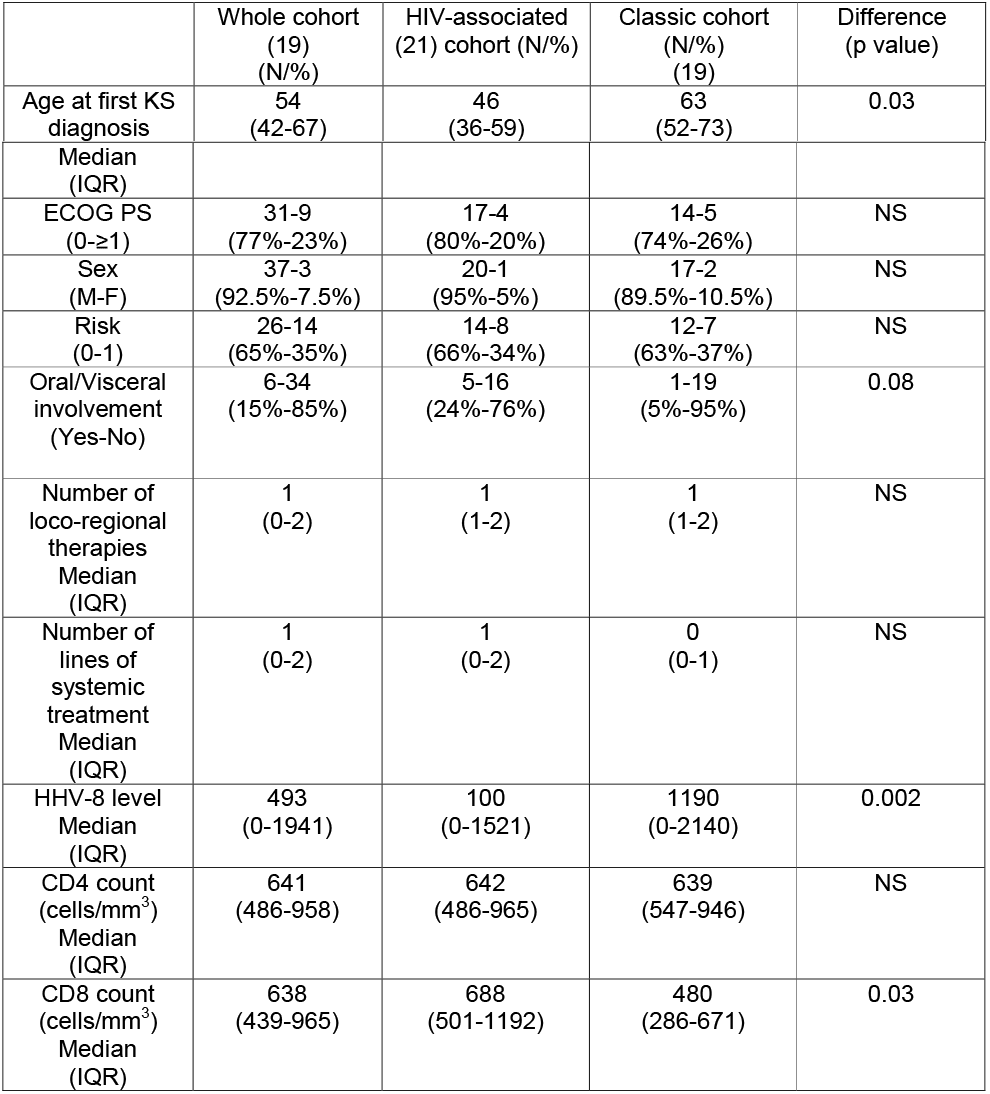
Baseline characteristics of the patient cohorts.

|  | Whole cohort<br>(19)<br>(N/%) | HIV-associated<br>(21) cohort (N/%) | Classic cohort<br>(N/%)<br>(19) | Difference<br>(p value) |
| --- | --- | --- | --- | --- |
| Age at first KS<br>diagnosis | 54<br>(42-67) | 46<br>(36-59) | 63<br>(52-73) | 0.03 |
| Median (IQR) |  |  |  |  |
| ECOG PS (0-≥1) | 31-9<br>(77%-23%) | 17-4<br>(80%-20%) | 14-5<br>(74%-26%) | NS |
| Sex (M-F) | 37-3<br>(92.5%-7.5%) | 20-1<br>(95%-5%) | 17-2<br>(89.5%-10.5%) | NS |
| Risk (0-1) | 26-14<br>(65%-35%) | 14-8<br>(66%-34%) | 12-7<br>(63%-37%) | NS |
| Oral/Visceral involvement (Yes-No) | 6-34<br>(15%-85%) | 5-16<br>(24%-76%) | 1-19<br>(5%-95%) | 0.08 |
| Number of loco-regional therapies<br>Median (IQR) | 1<br>(0-2) | 1<br>(1-2) | 1<br>(1-2) | NS |
| Number of lines of systemic treatment<br>Median (IQR) | 1<br>(0-2) | 1<br>(0-2) | 0<br>(0-1) | NS |
| HHV-8 level<br>Median (IQR) | 493<br>(0-1941) | 100<br>(0-1521) | 1190<br>(0-2140) | 0.002 |
| CD4 count (cells/mm<sup>3</sup>)<br>Median (IQR) | 641<br>(486-958) | 642<br>(486-965) | 639<br>(547-946) | NS |
| CD8 count (cells/mm<sup>3</sup>)<br>Median (IQR) | 638<br>(439-965) | 688<br>(501-1192) | 480<br>(286-671) | 0.03 |

### Targeted transcriptomic profiling reveals divergent polarisation of the tumour microenvironment in HIV-related versus classic KS

After removing samples with inadequate RNA quantity (N=7) and those failing quality check (N=8), 25 samples were retained for analysis: 13 in the HIV-associated cohort and 12 in the classic cohort. Differential gene set expression analysis showed significant upregulation of genes associated with innate and adaptive inflammatory responses in classic KS compared with HIV-associated cases (**Figure 1**). **Supplementary Table 5** reports the top 30 upregulated genes in the 2 cohorts. Pathway analysis demonstrated significant upregulation of gene transcriptional programmes related to allograft rejection, TNF-alpha, hypoxia, IL-6 and angiogenesis in classic KS, confirming enrichment of adaptive and innate immune pathways in classic cases (**Supplementary Figure 2A**). Conversely, transcripts related to glycolysis and epithelial to mesenchymal transition were upregulated in HIV-associated cases (**Supplementary Figure 2B**).

**Figure 2.**
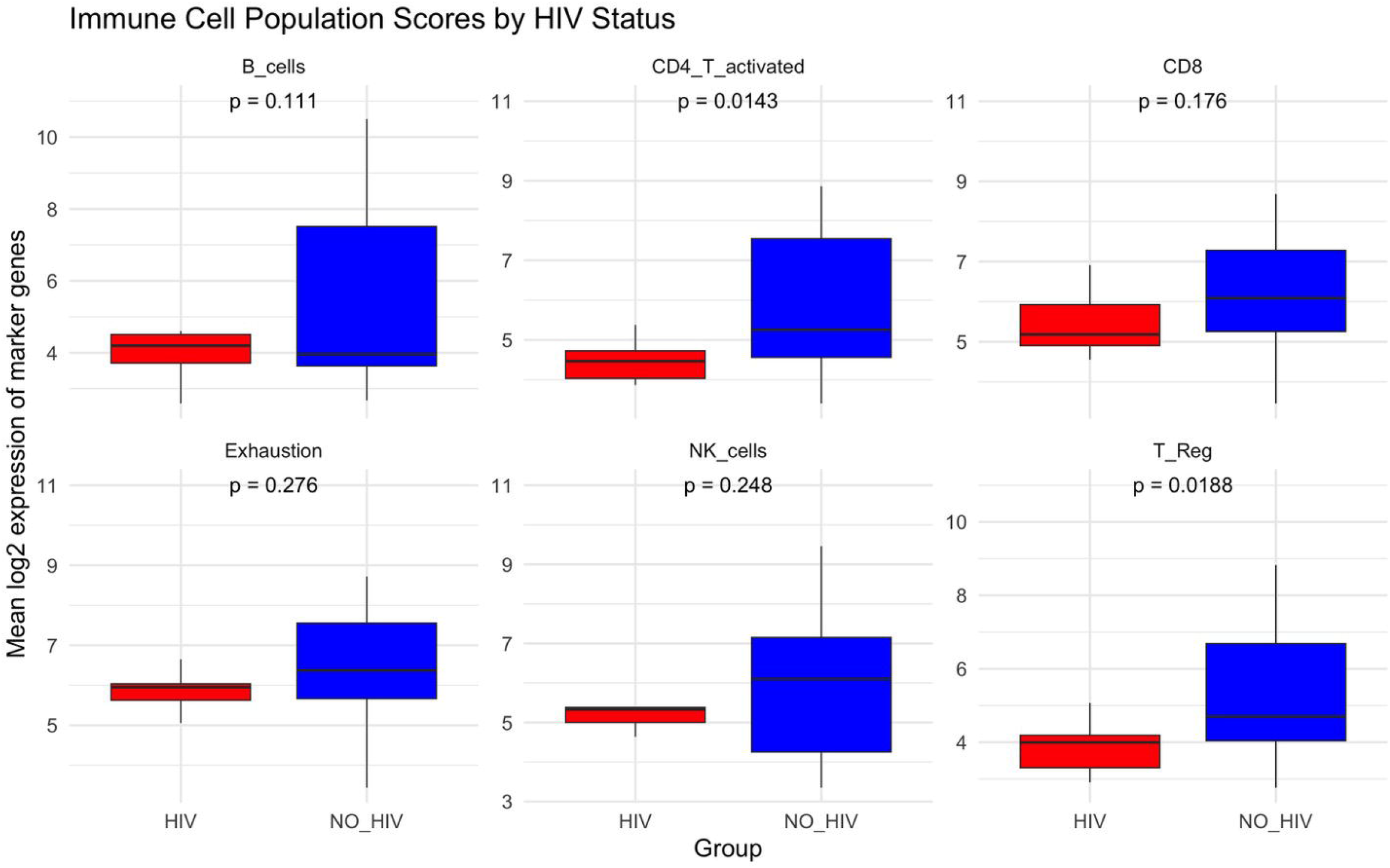
Comparison of immune cells population in HIV associated cases (red) and non-HIV cases (blue).

We then assessed the composition of the immune TME by calculating the immune score for 6 immune population as reported in the **Methods** section, and we observed higher representation of activated CD4 (p=0.031) and regulatory T-cells (p=0.019) in non-HIV cases, and a trend toward higher B-cell (p=0.11) and CD8 infiltration (p=0.18, **Figure 2**).

### Differential expression of intra-tumoral HHV-8 genes according to the HIV status

Because we found that plasma HHV-8 DNA concentration was significantly higher in classic KS cases, we hypothesised that HHV-8 replicative activity might be differentially regulated depending on HIV status. To prove this hypothesis, we tested for the differential expression of a panel of selected HHV-8 genes according to HIV co-infection status. We found a significantly higher expression of HHV-8 K8.1 and v-IRF1 in classic KS, even though it showed only a trend toward significance after FDR adjustment (FDR=0.085 and 0.124, respectively); only a modest correlation was observed between HHV-8 DNA levels and the intra-tumoral expression of selected HHV-8 genes (**Supplementary Figure 3**). We then calculated the HHV-8 score for each sample, defined as the mean log2 expression of the viral genes, and dichotomised the population based on the median HHV-8 score into “high” versus “low”. This confirmed a non-significant correlation between the HHV-8 score and plasma HHV-8 DNA. In the low HHV-8 group, the proportion of patients with HIV was numerically higher (66.7 % vs 33.3%), and similarly, the high HHV-8 group had a numerically higher proportion of non-HIV samples (61.5% vs 38.5%), suggesting that higher HHV-8 gene expression may be required for the progression of KS in the absence of HIV.

**Figure 3.**
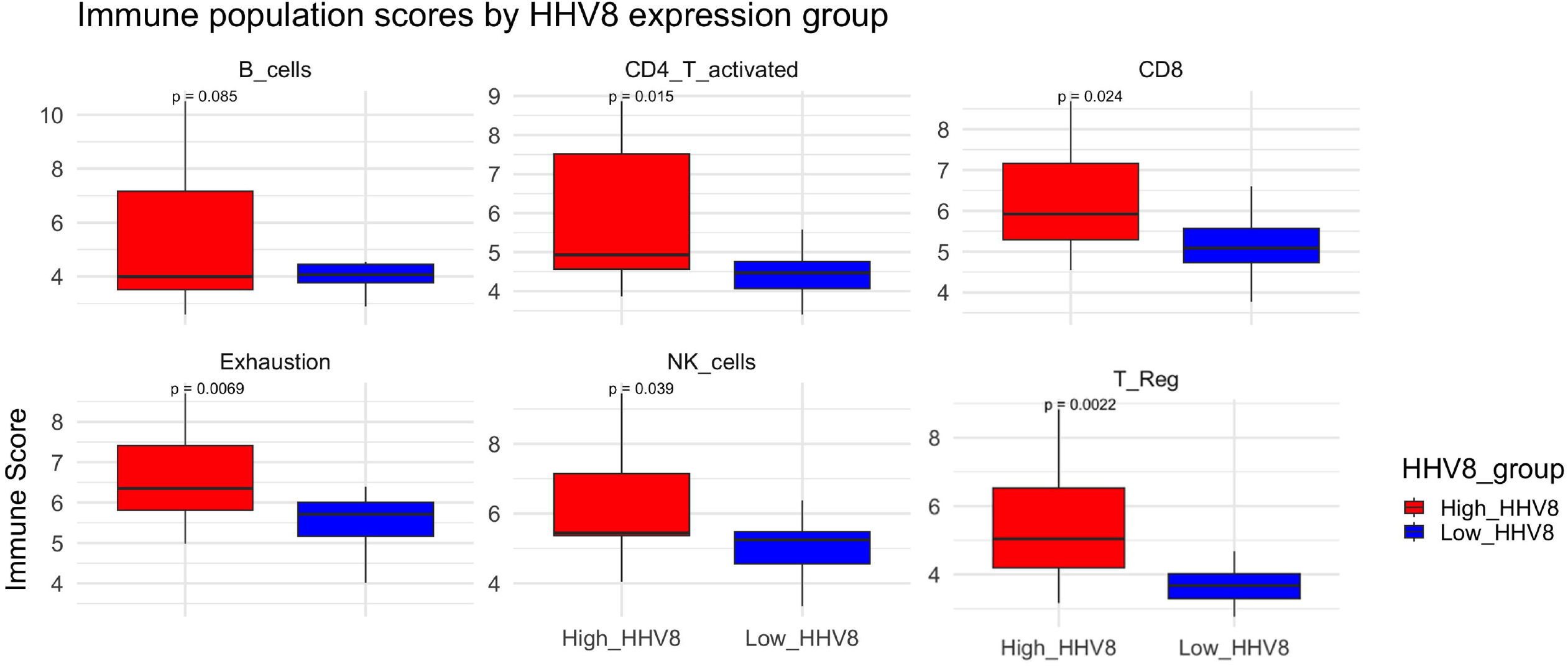
Comparison of immune cells population based on the HHV8 score. “High” is reported in red and “low” in blue.

### Intra-tumoral expression of HHV-8 genes is associated with distinctive composition of the tumour microenvironment

Subsequently, as numerically different HHV-8 gene expression was present in the two cohorts; to define the impact of HHV-8 on the TME, we assessed the correlation between the expression of HHV-8 genes in the tissue and of the human genes included in the panel. These revealed a positive correlation between the expression of the selected HHV-8 genes in the tumour and 202 host transcripts (**Supplementary Figure 4 and Supplementary Table 6**), the ORA showed a positive association between the expression of viral genes and inflammation pathways, as well as pathways related to cells proliferation and epithelial to mesenchymal transition (**Supplementary Figure 7**). As HHV-8 can exist in both latent and lytic phase in KS lesions, we assessed separately the expression of latent and lytic genes and their correlation TME transcripts. We observed that in the whole cohort, the mean expression of latent genes was higher compared to lytic ones (p=0.03), in accordance with latent phase considered as the predominant state of infection. Whilst most of the upregulated TME genes correlating with viral expression overlapped between latent and lytic genes (N=129), we identified a subset of genes that were exclusively associated with either lytic only (N=73) (**Supplementary Table 7**), or latent only (N=17) genes (**Supplementary Table 8**). The ORA pathway analysis showed that a stronger inflammatory response was associated with lytic genes whereas genes significantly associated only with latent genes contributed to tumour-associated pathways; however, none of the pathways remained significant after FDR adjustment. This suggests a possible differential effect of latent and lytic genes in influencing KS pathogenesis and progression.

**Figure 4.**
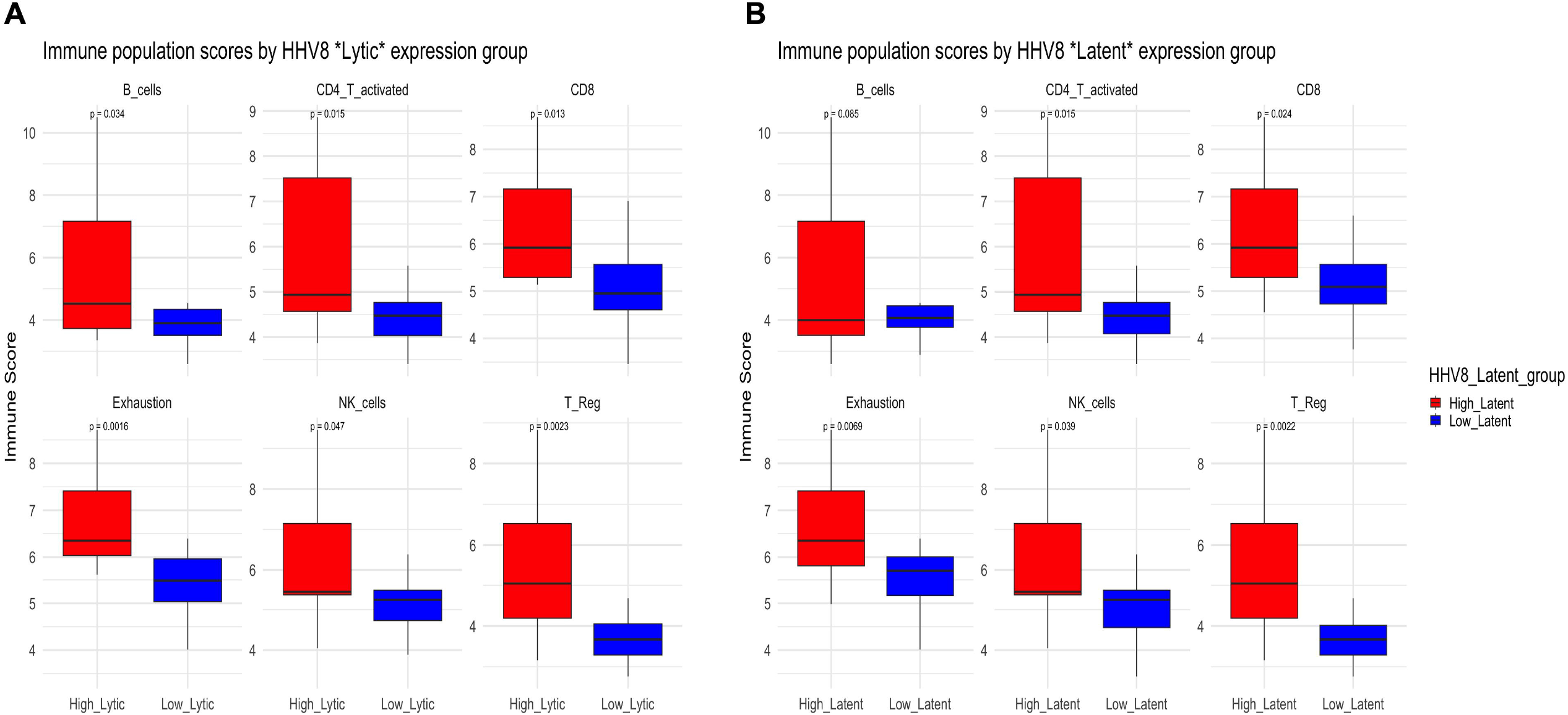
**A-B**. Comparison of immune cells population based on the Lytic (A) and Latent (B) HHV8 score. “High” is reported in red and “low” in blue.

We then compared the prevalence of each immune population, according to the HHV-8 score (**Figure 3**). It showed that high HHV-8 score was associated with higher expression of genes related to CD4 activated (p=0.013), CD8 activated (p=0.025) cells, NK (p=0.039), T-regulatory cells (p=0.002) and higher expression of gene sets associated with T-cell exhaustion (p=0.007). The differences were maintained when calculating separately the HHV-8 score for latent and lytic genes (**Figure 4A-B**).

### Multiplex immunofluorescence highlights differential distribution of T-cell subsets in HIV-associated KS

To validate the RNA sequencing data, we performed immune-fluorscence (IF) on 12 samples that had available material for both immune-fluorescence and RNA analysis (HIV=3; non-HIV =9). Based on the biological relevance and adjusted p value derived from the differential gene expression experiments across the two cohorts (**Table 2**), 12 protein markers of interest were selected to validate immune cell distribution across clinical groups (**Supplementary Tabls 3-4, Figure 5A-D**).

**Figure 5.**
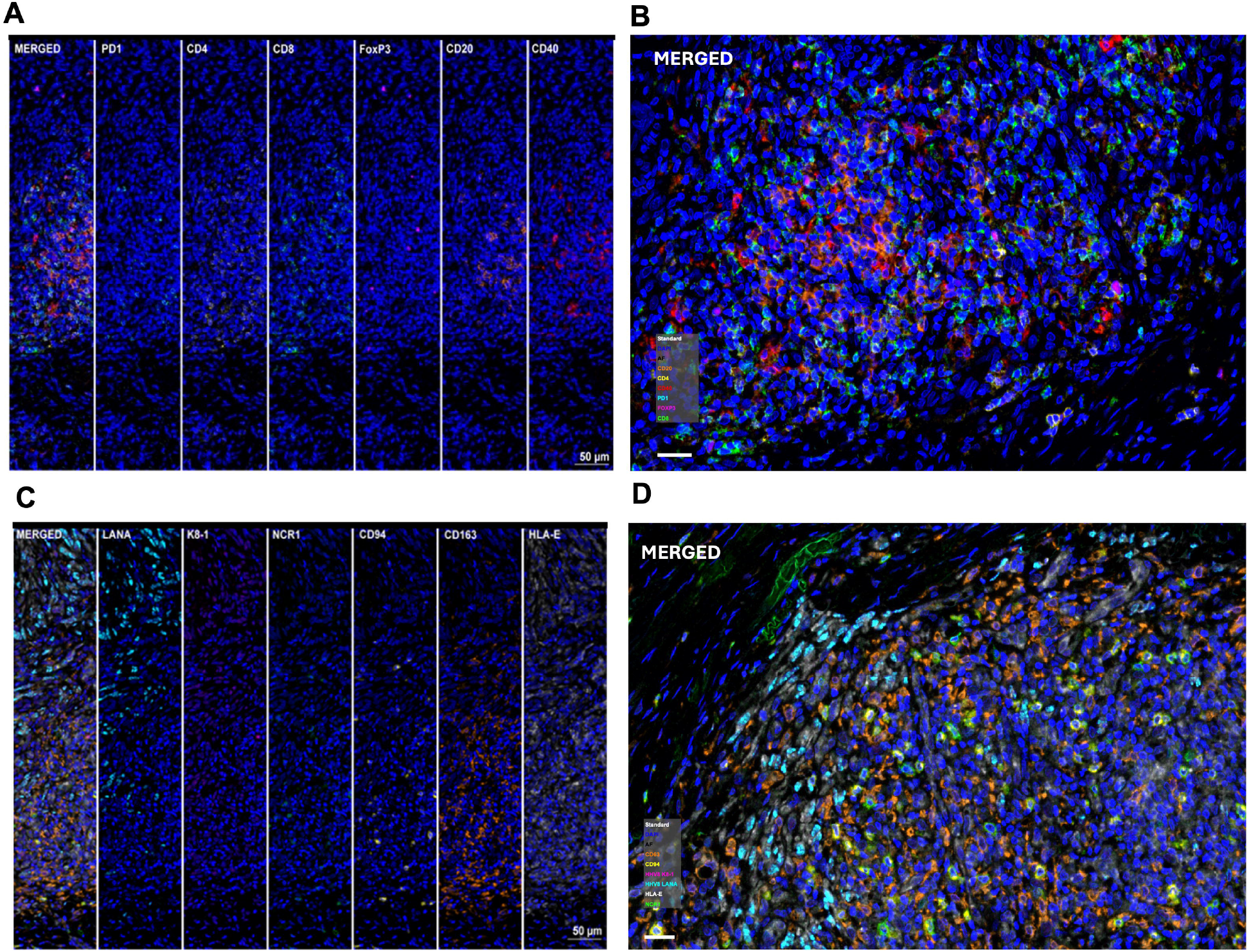
**A-B**. High-resolution image showing marker expression (Panel 1): Cyan, PD-1; yellow, CD4; green, CD8; purple, FoxP3; orange, CD20; red, CD40; blue, DAPI (nuclei); At 20x (A), scale bar at 50 µm, and 40x (B), scale bar at 20 µm. **Figure 5C-D**. High-resolution image showing marker expression (Panel 2): Cyan, LANA; purple, K8.1; green, NCR1; yellow, CD94; orange, CD163; white, HLA-E; blue, DAPI (nuclei). At 20x (C), scale bar at 50 µm, and 40x (D), scale bar at 20 µm.

**Table 2.** Target of antibodies used in the immunofluorescence panel, along with the adjusted p-value of the differential gene expression (HIV vs Non-HIV).

| Immunofluorescence marker | Adj.P. Value DEG |
| --- | --- |
| FOXP3 | 0.005 |
| CD4 | 0.05 |
| CD20 (*MS4A1*) | 0.32 |
| PD1 (*PDCD1*) | 0.22 |
| CD40 | 0.45 |
| CD8 | 0.89 |
| NCR1 | 0.26 |
| CD94 (*KLRD1*) | 0.35 |
| CD163 | 0.40 |
| HLA-E | 0.86 |
| HHV-8 K8.1 | 0.09 |
| HHV-8 LANA | 0.20 |

As reported in **Figure 6A-B**, we observed a higher proportion of CD4+ T-cell (p=0.05) and regulatory T-cells expressing FOXP3 (p=0.03) in the classic KS cohort, as well as a trend toward higher CD20+ B cells infiltration (p=0.06), confirming targeted transcriptomics findings. Similarly, no difference in the other cell subtypes was observed.

**Figure 6.**
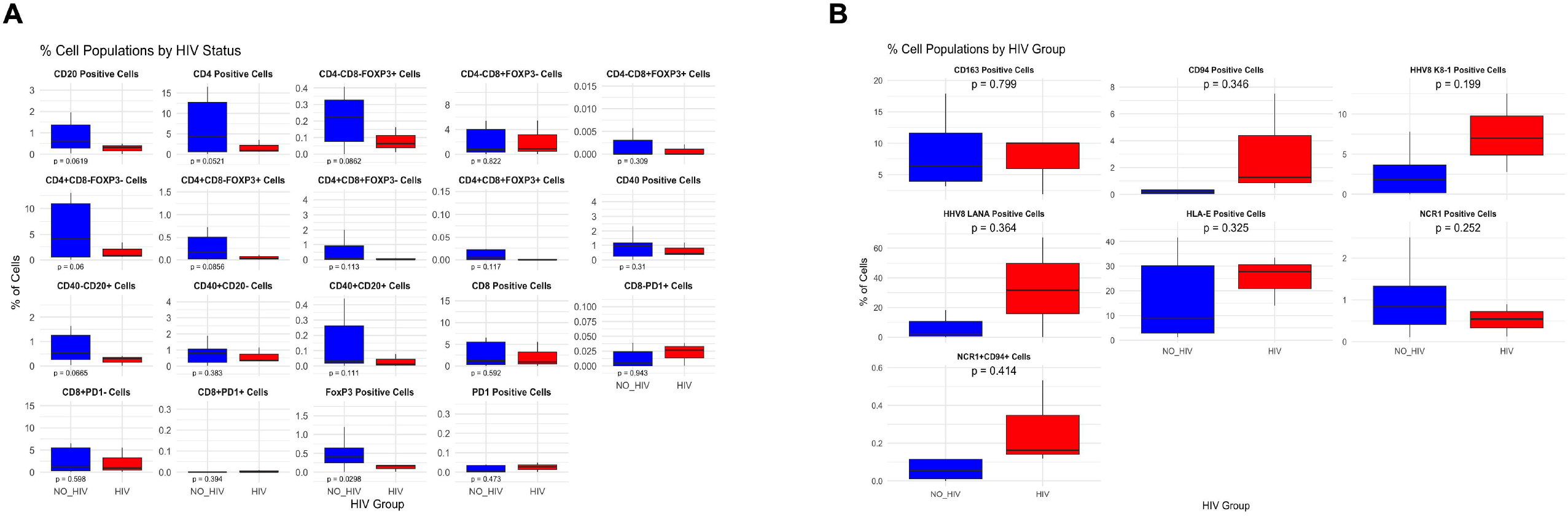
**A-B**. Comparison of immune cells population detected by immune-fluorescence based on the HIV status. No-HIV is reported in blue and HIV-associated is in red, in Panel 1 (A), and Panel 2 (B).

To assess the impact of HHV8 expression on the immune microenvironment at the protein level, IF cohort was stratified according to the percentage of LANA and K8.1 positive cells. The median percentage across all samples was calculated, and cases were classified as “high” or “low” based on whether their value was above or below the median. Samples classified as “high” for both LANA and K8.1 were defined as having “high HHV8 expression”, the remaining samples were classified as “low HHV8 expression”. The proportions of immune cells, defined by markers included in the same IF panel as LANA and K8.1, were compared between the two groups using Welch’s t-test. In total, five samples were categorized as “high HHV8” and seven as “low HHV8”. As showed in **Figure 7**, “High HHV8” was associated with significantly higher proportion of HLA-E positive cells (p=0.001), a trend toward higher proportion of NCR1+ NK cells (p=0.07), and higher proportion of NK cells expressing both NCR1 and CD94 (p=0.027), which upon binding with HLA-E inhibits NK activity. These findings confirm a direct role of HHV8 in harnessing the host immune response.

**Figure 7.**
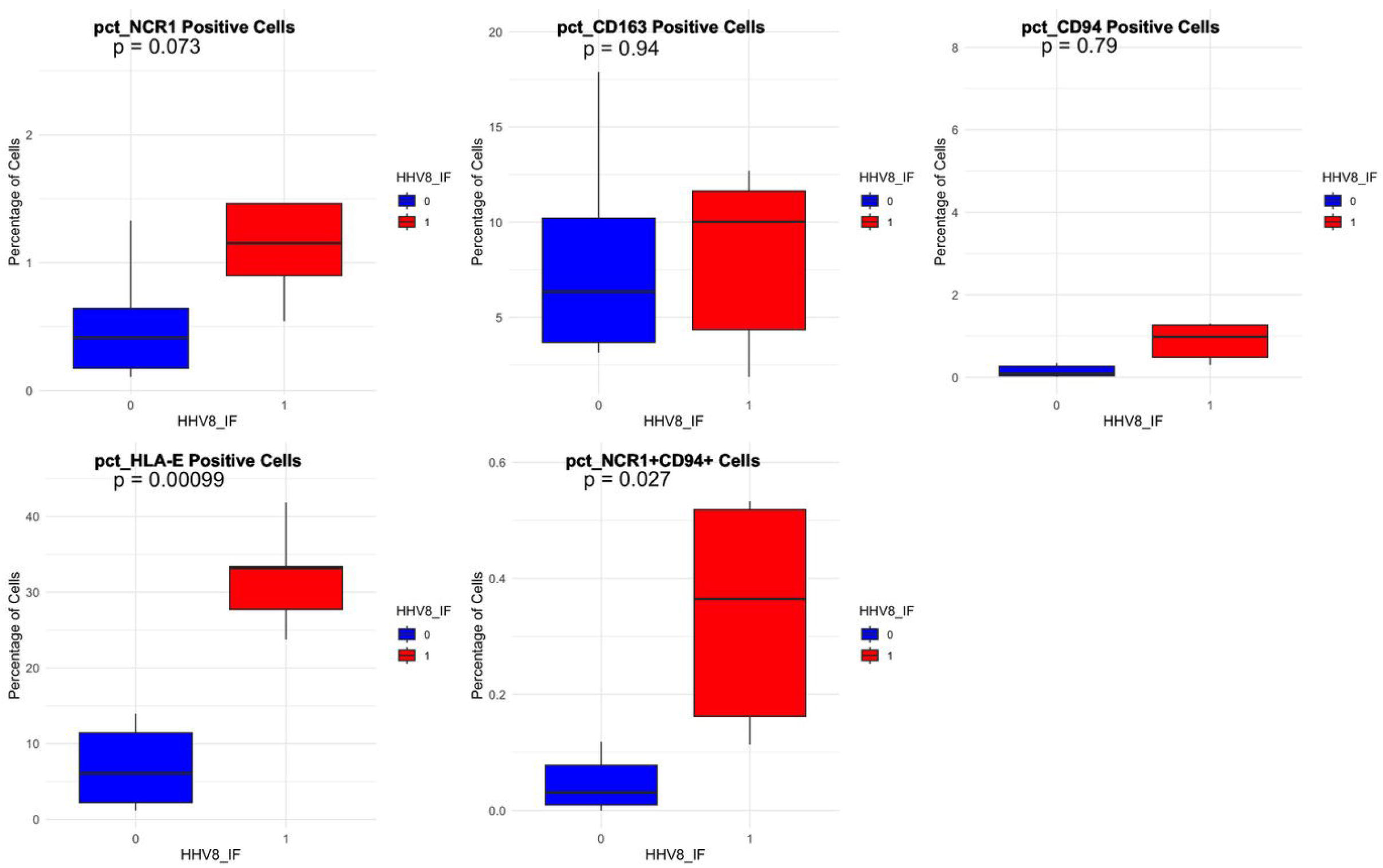
Comparison of immune cells population detected by immune-fluorescence based on the expression of LANA and K8.1 in immune-fluorescence in Panel 1. Low expression is reported in blue, high expression is reported in red.

## Discussion

HHV-8 infection is the unifying mechanism underlying the pathogenesis of the distinct classes of KS. By infecting blood and lymphatic vascular endothelial cells, HHV-8 directly contributes to tumorigenesis by promoting vascular proliferation, cell survival and apoptosis inhibition^2^. Alterations of adaptive immunity are fundamental pathogenic mechanisms that influence its development and progression. However, the functional characteristics of the KS tumour immune microenvironment occurring in the absence of clinically evident immune suppression are poorly defined, with detrimental implications for the development of novel therapies. In this study, we performed an in-depth characterisation of the tumour microenvironment of KS occurring in the absence of clinically significant immune suppression. Even if other studies have compared the TME of different subtypes of KS ^12 13^, this is the first study focusing on KS occurring in patients without clinically evident immune-suppression.

By comparing two KS cohorts, we show evidence of a more inflamed tumour immune microenvironment in classic KS compared to HIV-associated KS, despite comparable clinical characteristics. Our targeted RNA analysis using the Nanostring nCounter platform revealed an enrichment in inflammatory pathways, including TNFalpha and IL-6 in classic KS, up-regulation of pathways related to epithelial to mesenchymal transition and glycolysis was more common in HIV associated KS. This was paired with higher expression of genes ascribable to activated CD4 and regulatory T-cells in classic cases.

These results underscore different pathogenetic mechanisms between the two subtypes, which point towards the requirement of a more profound inflammatory dysregulation for the development of KS in the absence of HIV. These findings resonate with earlier evidence reported by Lidenge et al. in a comparative study, which was however confounded by the inclusion of AIDS-associated KS among the HIV-associated cases^14^.

Using multiplex immunofluorescence, we sought to validate the gene expression results, while also assessing the proportion of other cardinal cell populations involved in anti-tumour immune control. Our findings confirmed higher infiltration of CD4 and regulatory T-cells in classic KS cases, while observing a relative uniformity of the other immune populations of interest across the two cohorts.

As HHV-8 is intimately linked to KS pathogenesis, we investigated how transcriptional activation of key HHV-8 genes relates to the composition of the TME across the two cohorts. While most of the assessed HHV-8 genes did not correlate with the peripheral HHV-8 DNA levels, a positive correlation was found between HHV-8 DNA level and intra-tumoral *ORF74* and *ORF75* - encoding two lytic proteins. Furthermore, a trend toward higher expression of *vIRF* and *K8*.*1* was observed in classic KS.

These results are provocative in suggesting the presence of a more pronounced viral lytic replication for KS pathogenesis in the absence of HIV. However, the limited sample size and lack of skin biopsies from non-tumoral sites constrain the interpretability of these findings. In our study HHV-8 gene expression was positively associated with the overactivation of several inflammatory cellular pathways, including IL-6, TNFα, IFNγ and KS specific pathways such as endothelial to mesenchymal transition^15-17^ and angiogenesis independently from the presence of HIV. These results are consistent with the known complex interaction between HHV-8 and host immunity. HHV-8 is known to downregulate the adaptive immune response to avoid epitope-specific immune clearance. On the other hand, the virus can directly activate a pro-tumorigenic inflammatory response, which is in turn necessary for KS development^18^.

In particular, it can activate the IL-6 pathway by its viral homologue^19^, as well as inducing endothelial to mesenchymal transition and angiogenesis through among others, *LANA* and *vFLIP* expression^20^. Despite being known to inhibit the host response, the production of IFN gamma and TNF alpha is pivotal for KS development, and can be directly stimulated by viral proteins, as OX2^2^. Interestingly, we also observed a higher degree of T-cell infiltration and immune exhaustion in the presence of high HHV-8 latent or lytic transcriptional activity, independently from the HIV status. This highlights the central role of virus-host interactions as a tumourigenic mechanism in KS and a potential source of targeted treatment strategies in this oncological indication.

We acknowledge several limitations that might hamper the interpretation of our results. First, the retrospective nature of the study and the use of archival biological samples significantly limited the sample size. Second, the only partial overlap between RNA and immunofluorescence datasets restricted our power to validate the findings at the protein level. Third, the absence of multiple sampling from the same patients, as well as the lack of samples from normal skin or from patients affected by other KS subtypes, limits both the interpretation and generalisation of our data. Despite these limitations, our study benefits from the use of two matched cohorts of KS occurring in the absence of immune suppression, providing a comprehensive comparative evaluation of two distinct clinical subtypes.

Overall, our data indicate that—despite similar clinical characteristics—a more pronounced activation of both adaptive and innate immunity pathways is evident in classic KS compared to HIV-associated KS occurring in the absence of immune suppression, despite an overall similar phenotypic composition of the tumour immune infiltrate. This difference may reflect distinct biological mechanisms that warrant dedicated preclinical investigation. We also highlighted the pivotal role of intra-tumoral HHV-8 gene transcription in shaping the local inflammatory microenvironment, providing a rationale to harness anti-viral immunity in the development of novel anti-KS strategies.

## Acknowledgements

The authors would like to thank the ECMC laboratory at Hammersmith Hospital (Naina Patel), the NIHR Imperial Experimental Cancer Medicine Centre, the Imperial College BRC, and Imperial College Healthcare Tissue Bank for the infrastructural support and the sample processing.

## Notes

**Conflict of interest:** CAMF received lecture fees from AstraZeneca and Eisai, and consultancy fees from MAIA biotechnology. MB received lecture fees from ViiV and Gilead Sciences, and institutional grants from Gilead Sciences. DJP received consulting fees from Mina Therapeutics, EISAI, Roche, Avamune, DaVolterra, Mursla, H3B, Ipsen, Boston Scientific, Starpharma, Exact Sciences, AstraZeneca, and LIfT Biosciences, and has received honoraria from Roche, BMS, EISAI, and Boston Scientific, and travel support from Roche, BMS, and MSD; received institutional grants from BMS, MSD, and GSK. All other authors declare no competing interests.

### Competing Interest Statement

CAMF received lecture fees from AstraZeneca and Eisai, and consultancy fees from MAIA biotechnology. MB received lecture fees from ViiV and Gilead Sciences, and institutional grants from Gilead Sciences. DJP received consulting fees from Mina Therapeutics, EISAI, Roche, Avamune, DaVolterra, Mursla, H3B, Ipsen, Boston Scientific, Starpharma, Exact Sciences, AstraZeneca, and LIfT Biosciences, and has received honoraria from Roche, BMS, EISAI, and Boston Scientific, and travel support from Roche, BMS, and MSD; received institutional grants from BMS, MSD, and GSK. All other authors declare no competing interests.

