## Supplementary material for "In depth characterisation of the tumour microenvironment reveals HHV-8 dependent immune regulation in HIV-associated and classic Kaposi Sarcoma": Supplementary.docx

*Fulgenzi CAM et al.*

**Supplementary Materials.**

**Table of Contents**

**SUPPLEMENTARY METHODS.**

**Supplementary Method 1.** Gene expression data processing and normalization. **Pag.2**

**Supplementary Method 2**. Immune fluorescence. **Pag.2**

**SUPPLEMENTARY Figures.**

**Supplementary Figure 1A-B**. Progression free survival in the clinical cohort. **Pag.3**

**Supplementary Figure 2A-B.** Pathway analysis of the upregulated genes in each cohort. **Pag.3**

**Supplementary Figure 3.** Correlation between the peripheral HHV-8 DNA and

the intra-tumoral expression of HHV-8 genes	**Pag.3**

**Supplementary Figure 4.** Correlation between the expression of host and viral genes. **Pag.4**

**Supplementary Figure 5.** Pathway analysis of the genes positively associated with

HHV8 expression	**Pag.4**

**SUPPLEMENTARY Tables.**

**Supplementary Table 1.** PanCancer IO 360 Panel. **Pag.5-36**

**Supplementary Table 2.** HHV8 genes included in the Nanostring nCounter analysis. **Pag. 36**

**Supplementary Tables 3-4.** List of Antibodies and BOND working conditions used for IHC

and mIF panels. **Pag.37**

**Supplementary Tables 5.** Top 30 genes differentially expressed in the 2 cohorts. **Pag.37-38**

**Supplementary Tables 6.** Host genes associated with HHV8 genes. **Pag. 38-43**

**Supplementary Tables 7.** Host genes associated with HHV8 lytic genes. **Pag. 43-44**

**Supplementary Tables 8.** Host genes associated with HHV8 latent genes. **Pag. 44-45**

**Supplementary Method 1.** *Gene expression data processing and normalization.*

Raw RCC files were downloaded and processed with nSolver Analysis Software (NanoString Technologies). After removing samples that did not meet the quality check standards, normalization was performed by background correction using the geometric mean of negative controls, followed by positive control normalization using the geometric mean of internal spike-in controls and housekeeping gene normalization (geometric mean), retaining only genes with mean counts >100 and coefficient of variation within ±10%. The following housekeeping (HK) genes were retained and used for normalization: PSMC4, ABCF1, PUM1, TMUB2, SF3A1, OAZ1. Downstream analysis was performed in R. Normalized expression values were log2-transformed prior to analysis. Differential gene expression (DEG) analysis between the 2 groups was performed using linear models with empirical Bayes moderation (via limma package in R). Hierarchical clustering and heatmaps were generated using Euclidean distance and complete linkage. P values were adjusted for multiple comparisons using the Benjamini-Hochberg methos. An adjusted p value <0.05 and log2 fold change>0.5 were used to determine significance.

Genes with P < 0.05 and log2FC > 0.5 were considered differentially expressed and used for Over-Representation Analysis (ORA). Two separate ORA analyses were performed using the enricher function from the clusterProfiler R package with MSigDB Hallmark gene sets. Enrichment of biological processes was assessed independently for each gene set group.

Significance was determined using a hypergeometric test, followed by multiple testing correction (Benjamini-Hochberg). The top enriched pathways were visualized using dot plots. Directional analysis allowed the identification of pathways specifically enriched in each group.

The abundance of immune cells population was inferred from the Nanostring Pancancer360IO panel by calculating the mean expression of predefined marker gene corresponding to 5 distinct immune population. For each sample, log2-transformed normalised expression values were extracted and averaged for gene signatures specific to CD8⁺ activated T cells, CD4⁺ activated T cells, natural killer (NK) cells, B cells, and immune checkpoint molecules. As already published11, the CD8⁺ T cell gene set comprised GZMB, CD8A, PRF1, and CD8B; CD4⁺ activated T cells were represented by CD4, IL26 and IL17A; NK cells from the expression of NCR1, KLRB1, KLRC1 and KLRD; T-regulatory from FOXP3, IL2RA (CD25), CTLA4, and GITR; B cells from BLK, CD19, and MS4A1. Finally, immune checkpoint molecules expression was assessed using HAVCR2 (TIM-3), LAG3, CTLA4, PDCD1 (PD-1), and TIGIT. For each immune population, an immune score was calculated as the mean log2 expression of the respective marker genes detected in the Nanostring panel and compared between HIV and Non-HIV cases using unpaired two-sided Student’s t-tests

To investigate the impact of viral transcriptional activity, we quantified the expression of the selected HHV-8 genes, including K12, K8.1, ORF16, ORF50, ORF71, ORF72, ORF73, ORF74, ORF75, and v-IRF1. For each sample, a viral HHV-8 score was computed as the mean log2 expression of these viral genes. Samples were then stratified into “HHV-8-High” and “HHV-8-Low” groups based on the median viral score across all cases.

Immune population scores were subsequently compared between the High HHV-8 and Low HHV-8 groups using unpaired two-sided Student’s t-tests.

All statistical analyses were performed in R (v4.x) using the limma, dplyr, ggplot2, and ggpubr packages.

**Supplementary Method 2.** *.*

Initially, single-marker IHC assays were conducted on FFPE sections to individually optimize conditions for each target. This included determining appropriate heat-induced epitope retrieval (HIER) settings, antibody incubation durations, optimal concentrations, and assessing whether signal amplification was required. Details on the antibodies and specific staining parameters are summarized in Table (**Supplementary Table 3**).

To determine the optimal sequence for marker application in each multiplex panel, control slides were subjected to a series of single IHC tests with modifications in the number of HIER cycles. This optimization was repeated across all primary antibodies intended for mIF use, ensuring the most effective pairing and order for antibody and Opal combinations.

For multiplex staining, FFPE sections were deparaffinized, rehydrated, and underwent HIER using the BOND RX platform. Primary antibody incubation was standardized to 30 minutes. Detection employed either Opal Polymer HRP secondary reagents or SignalStain® Boost IHC Detection RAT (Cat. 72838S, Cell Signaling Technology, NL), followed by deposition of the corresponding Opal fluorophores from the Opal 7-Plex kit (Akoya Biosciences, Menlo Park, CA, USA). Each staining round was followed by an additional HIER step to remove antibody complexes, preparing the tissue for subsequent marker application. After completing all rounds, DAPI was added for nuclear counterstaining, and slides were mounted using X6 Immu-Mount (Fisher Scientific, UK) and cured at room temperature in the dark for one hour. Imaging was performed using the Vectra Polaris scanner (Akoya Biosciences), using the PhenoImager HT 2.0 software.

Digital image analysis was performed using the HALO Image Analysis Platform (version 3.6.4134, Indica Labs, Inc.). The mIF images were processed by IPU trained pathology analysts using the Highplex FL v4.2.14 module for segmentation, annotation, phenotyping, and quantification. Tumor areas were manually annotated, excluding necrotic zones and technical artifacts such as folds or detached fragments. Cell segmentation thresholds were tailored to the morphological features of each sample to ensure representativeness. Marker-specific signal intensity and staining patterns were reviewed to fine-tune positivity thresholds for phenotyping, based on controls. All relevant cellular phenotypes were defined through specific combinations of marker expression. Quantification was then carried out using these parameters, and results were reported as the number of positive cells per unit tissue area (μm²).

**Supplementary Figure 1A-B.** Kaplan Maier plots reporting the progression free survival in the whole cohort (A) and stratified by HIV status (B).

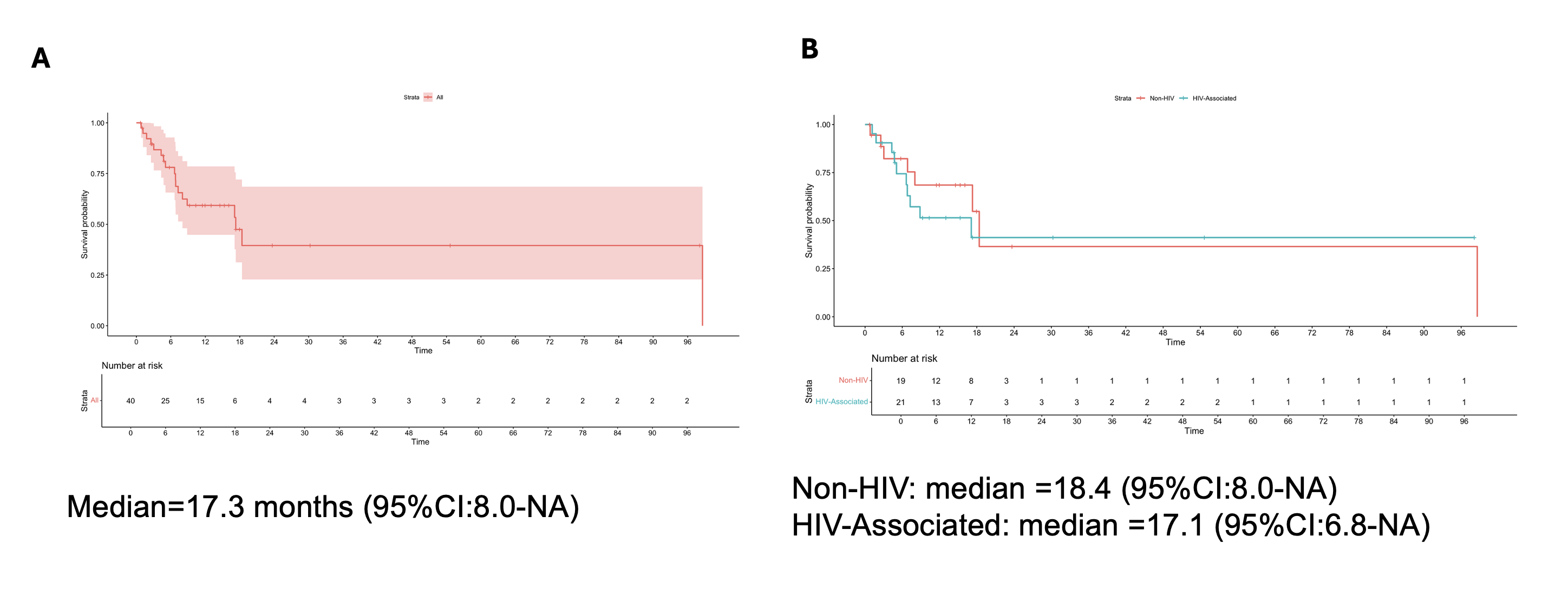

**Supplementary Figure 2A-B**. Over-representation analysis (ORA) showing upregulated pathways in Non-HIV (A) and HIV cases.

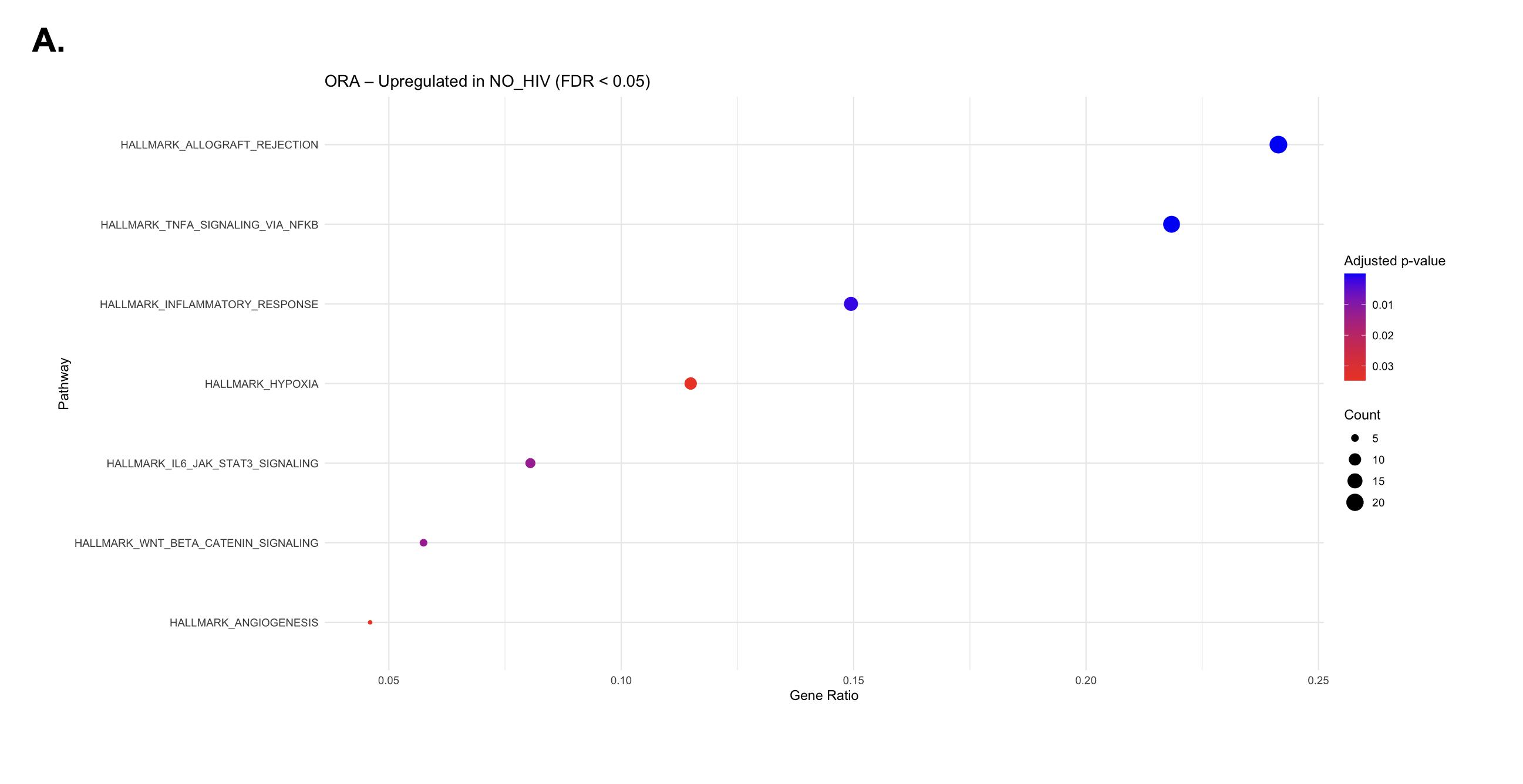

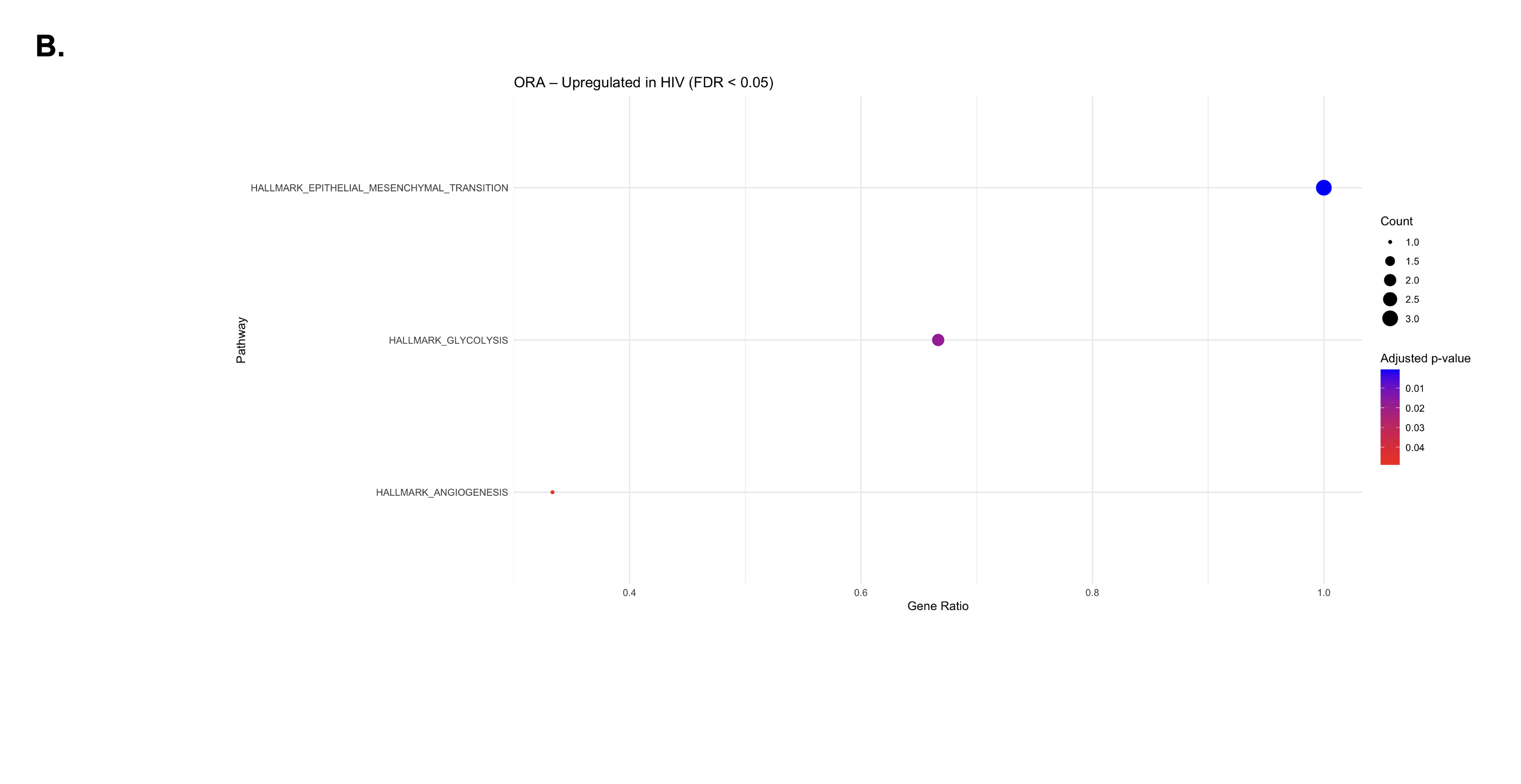

**Supplementary Figure 3.** Spearman correlation between HHV-8 DNA level and the intra-tumoral expression of HHV-8 genes

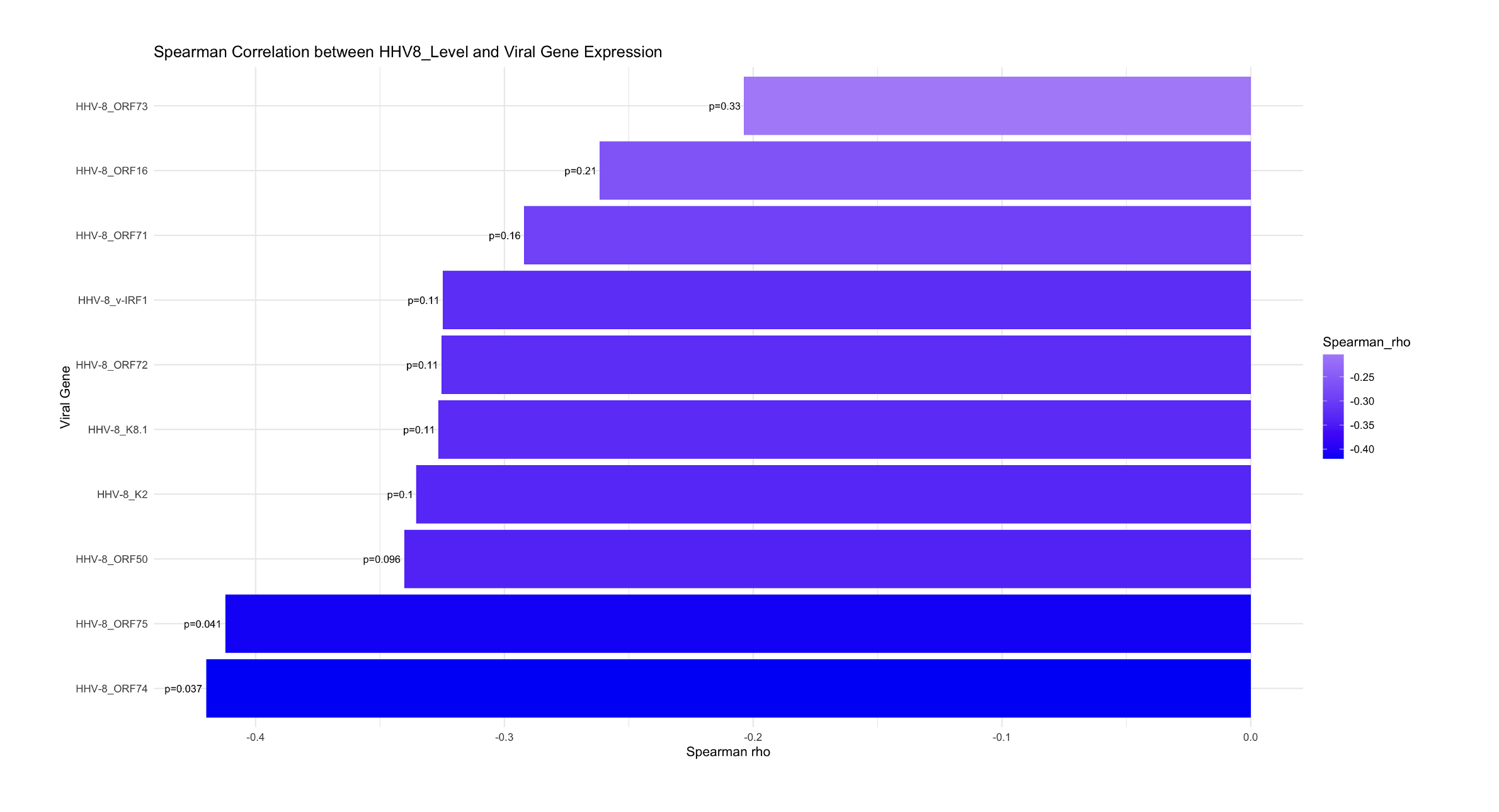

**Supplementary Figure 4.** Top 50 cellular genes (selected by FDR) significantly associated with the expression of HHV-8 genes**.**

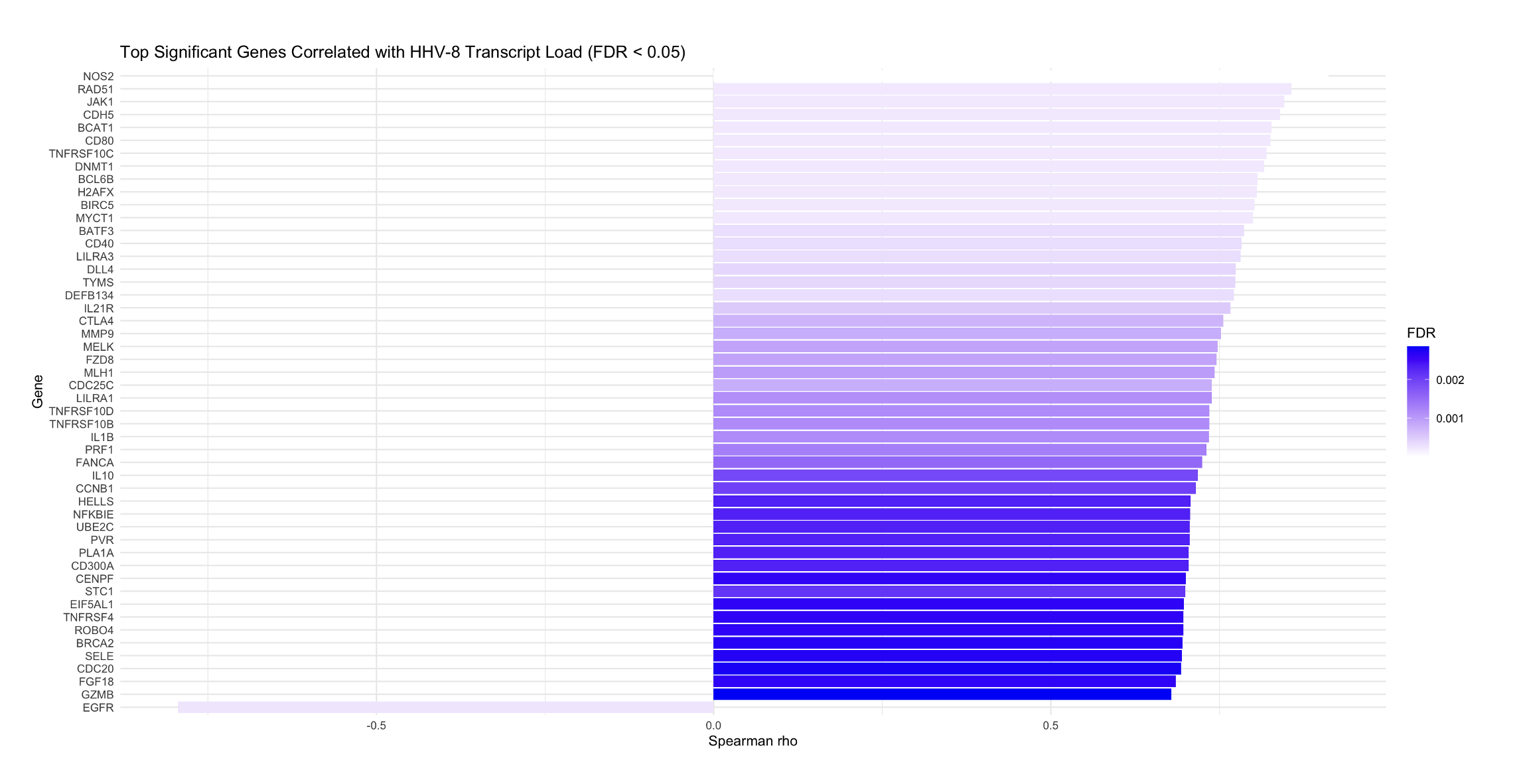

**Supplementary Figure 5.** Over-representation analysis (ORA) showing the host pathways that are positively associated with the expression of HHV-8 genes.

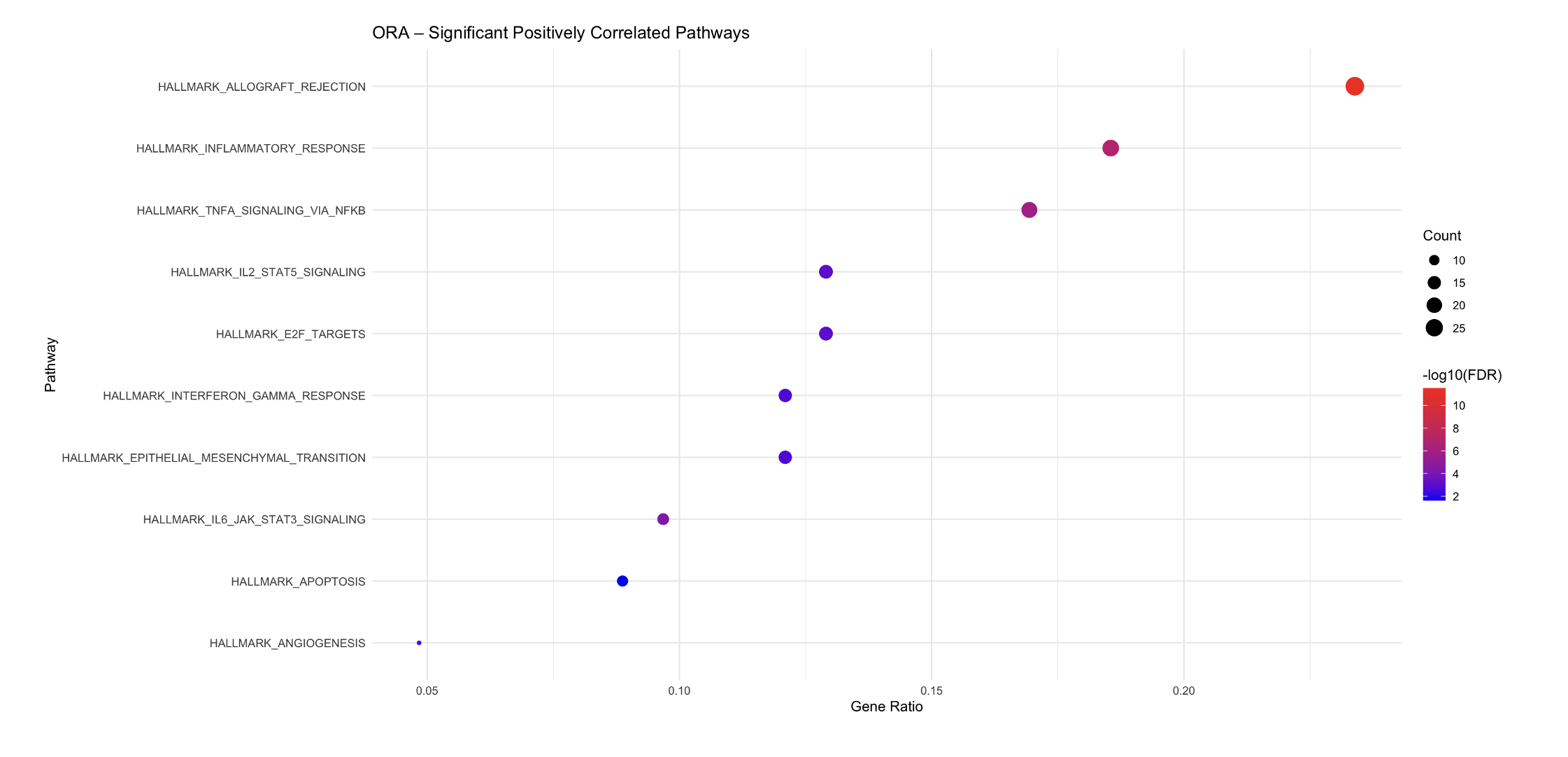

**Supplementary Table 1.** Complete list of the targeted genes profiled using the NanoString 360 PanCancer IO panel.

| **Gene Name** | **Official Full Name** |
| --- | --- |
| A2M | alpha-2-macroglobulin |
| ABCB1 | ATP-binding cassette, sub-family B (MDR/TAP), member 1 |
| ABL1 | c-abl oncogene 1, non-receptor tyrosine kinase |
| ADA | adenosine deaminase |
| ADORA2A | adenosine A2a receptor |
| AICDA | activation-induced cytidine deaminase |
| AIRE | autoimmune regulator |
| AKT3 | v-akt murine thymoma viral oncogene homolog 3 (protein kinase B, gamma) |
| ALCAM | activated leukocyte cell adhesion molecule |
| AMBP | alpha-1-microglobulin/bikunin precursor |
| AMICA1 | adhesion molecule, interacts with CXADR antigen 1 |
| ANP32B | acidic (leucine-rich) nuclear phosphoprotein 32 family, member B |
| ANXA1 | annexin A1 |
| APOE | apolipoprotein E |
| APP | amyloid beta (A4) precursor protein |
| ARG1 | arginase, liver |
| ARG2 | arginase, type II |
| ATF1 | activating transcription factor 1 |
| ATF2 | activating transcription factor 2 |
| ATG10 | autophagy related 10 |
| ATG12 | autophagy related 12 |

| ATG16L1 | autophagy related 16-like 1 (S.  cerevisiae) |
| --- | --- |
| ATG5 | autophagy related 5 |
| ATG7 | autophagy related 7 |
| ATM | ataxia telangiectasia mutated |
| AXL | AXL receptor tyrosine kinase |
| BAGE | B melanoma antigen |
| BATF | basic leucine zipper transcription factor, ATF-like |
| BAX | BCL2-associated X protein |
| BCL10 | B-cell CLL/lymphoma 10 |
| BCL2 | B-cell CLL/lymphoma 2 |
| BCL2L1 | BCL2-like 1 |
| BCL6 | B-cell CLL/lymphoma 6 |
| BID | BH3 interacting domain death agonist |
| BIRC5 | baculoviral IAP repeat containing 5 |
| BLK | B lymphoid tyrosine kinase |
| BLNK | B-cell linker |
| BMI1 | BMI1 polycomb ring finger oncogene |
| BST1 | bone marrow stromal cell antigen 1 |
| BST2 | bone marrow stromal cell antigen 2 |
| BTK | Bruton agammaglobulinemia tyrosine kinase |
| BTLA | B and T lymphocyte associated |
| C1QA | complement component 1, q subcomponent, A chain |
| C1QB | complement component 1, q subcomponent, B chain |
| C1QBP | complement component 1, q subcomponent binding protein |
| C1R | complement component 1, r subcomponent |

| C1S | complement component 1, s subcomponent |
| --- | --- |
| C2 | complement component 2 |
| C3 | complement component 3 |
| C3AR1 | complement component 3a receptor 1 |
| C4B | complement component 4B (Chido blood group) |
| C4BPA | complement component 4 binding protein, alpha |
| C5 | complement component 5 |
| C6 | complement component 6 |
| C7 | complement component 7 |
| C8A | complement component 8, alpha polypeptide |
| C8B | complement component 8, beta polypeptide |
| C8G | complement component 8, gamma polypeptide |
| C9 | complement component 9 |
| CAMP | cathelicidin antimicrobial peptide |
| CARD11 | caspase recruitment domain family, member 11 |
| CARD9 | caspase recruitment domain family, member 9 |
| CASP1 | caspase 1, apoptosis-related cysteine peptidase |
| CASP10 | caspase 10, apoptosis-related cysteine peptidase |
| CASP3 | caspase 3, apoptosis-related cysteine peptidase |
| CASP8 | caspase 8, apoptosis-related cysteine peptidase |
| CCL1 | chemokine (C-C motif) ligand 1 |
| CCL11 | chemokine (C-C motif) ligand 11 |
| CCL13 | chemokine (C-C motif) ligand 13 |
| CCL14 | chemokine (C-C motif) ligand 14 |
| CCL15 | chemokine (C-C motif) ligand 15 |

| CCL16 | chemokine (C-C motif) ligand 16 |
| --- | --- |
| CCL17 | chemokine (C-C motif) ligand 17 |
| CCL18 | chemokine (C-C motif) ligand 18 (pulmonary and activationregulated) |
| CCL19 | chemokine (C-C motif) ligand 19 |
| CCL2 | chemokine (C-C motif) ligand 2 |
| CCL20 | chemokine (C-C motif) ligand 20 |
| CCL21 | chemokine (C-C motif) ligand 21 |
| CCL22 | chemokine (C-C motif) ligand 22 |
| CCL23 | chemokine (C-C motif) ligand 23 |
| CCL24 | chemokine (C-C motif) ligand 24 |
| CCL25 | chemokine (C-C motif) ligand 25 |
| CCL26 | chemokine (C-C motif) ligand 26 |
| CCL27 | chemokine (C-C motif) ligand 27 |
| CCL28 | chemokine (C-C motif) ligand 28 |
| CCL3 | chemokine (C-C motif) ligand 3 |
| CCL3L1 | chemokine (C-C motif) ligand 3-like 1 |
| CCL4 | chemokine (C-C motif) ligand 4 |
| CCL5 | chemokine (C-C motif) ligand 5 |
| CCL7 | chemokine (C-C motif) ligand 7 |
| CCL8 | chemokine (C-C motif) ligand 8 |
| CCND3 | cyclin D3 |
| CCR1 | chemokine (C-C motif) receptor 1 |
| CCR2 | chemokine (C-C motif) receptor 2 |
| CCR3 | chemokine (C-C motif) receptor 3 |
| CCR4 | chemokine (C-C motif) receptor 4 |

| CCR5 | chemokine (C-C motif) receptor 5 (gene/pseudogene) |
| --- | --- |
| CCR6 | chemokine (C-C motif) receptor 6 |
| CCR7 | chemokine (C-C motif) receptor 7 |
| CCR9 | chemokine (C-C motif) receptor 9 |
| CCRL2 | chemokine (C-C motif) receptor-like 2 |
| CD14 | CD14 molecule |
| CD160 | CD160 molecule |
| CD163 | CD163 molecule |
| CD164 | CD164 molecule, sialomucin |
| CD180 | CD180 molecule |
| CD19 | CD19 molecule |
| CD1A | CD1a molecule |
| CD1B | CD1b molecule |
| CD1C | CD1c molecule |
| CD1D | CD1d molecule |
| CD1E | CD1e molecule |
| CD2 | CD2 molecule |
| CD200 | CD200 molecule |
| CD207 | CD207 molecule, langerin |
| CD209 | CD209 molecule |
| CD22 | CD22 molecule |
| CD24 | CD24 molecule |
| CD244 | CD244 molecule, natural killer cell receptor 2B4 |
| CD247 | CD247 molecule |
| CD27 | CD27 molecule |
| CD274 | CD274 molecule |

| CD276 | CD276 molecule |
| --- | --- |
| CD28 | CD28 molecule |
| CD33 | CD33 molecule |
| CD34 | CD34 molecule |
| CD36 | CD36 molecule (thrombospondin receptor) |
| CD37 | CD37 molecule |
| CD38 | CD38 molecule |
| CD3D | CD3d molecule, delta (CD3-TCR complex) |
| CD3E | CD3e molecule, epsilon (CD3-TCR complex) |
| CD3EAP | CD3e molecule, epsilon associated protein |
| CD3G | CD3g molecule, gamma (CD3-TCR complex) |
| CD4 | CD4 molecule |
| CD40 | CD40 molecule, TNF receptor superfamily member 5 |
| CD40LG | CD40 ligand |
| CD44 | CD44 molecule (Indian blood group) |
| CD46 | CD46 molecule, complement regulatory protein |
| CD47 | CD47 molecule |
| CD48 | CD48 molecule |
| CD5 | CD5 molecule |
| CD53 | CD53 molecule |
| CD55 |  |
| CD58 | CD58 molecule |
| CD59 | CD59 molecule, complement regulatory protein |
| CD6 | CD6 molecule |
| CD63 | CD63 molecule |

| CD68 | CD68 molecule |
| --- | --- |
| CD7 | CD7 molecule |
| CD70 | CD70 molecule |
| CD74 |  |
| CD79A | CD79a molecule, immunoglobulinassociated alpha |
| CD79B | CD79b molecule, immunoglobulinassociated beta |
| CD80 | CD80 molecule |
| CD81 | CD81 molecule |
| CD83 | CD83 molecule |
| CD84 | CD84 molecule |
| CD86 | CD86 molecule |
| CD8A | CD8a molecule |
| CD8B | CD8b molecule |
| CD9 | CD9 molecule |
| CD96 | CD96 molecule |
| CD97 | CD97 molecule |
| CD99 | CD99 molecule |
| CDH1 | cadherin 1, type 1, E-cadherin (epithelial) |
| CDH5 | cadherin 5, type 2 (vascular endothelium) |
| CDK1 | cyclin-dependent kinase 1 |
| CDKN1A | cyclin-dependent kinase inhibitor 1A (p21, Cip1) |
| CEACAM1 | carcinoembryonic antigen-related cell adhesion molecule 1 (biliary glycoprotein) |

| CEACAM6 | carcinoembryonic antigen-related cell adhesion molecule 6 (nonspecific cross reacting antigen) |
| --- | --- |
| CEACAM8 | carcinoembryonic antigen-related cell adhesion molecule 8 |
| CEBPB | CCAAT/enhancer binding protein (C/EBP), beta |
| CFB | complement factor B |
| CFD | complement factor D (adipsin) |
| CFI | complement factor I |
| CFP | complement factor properdin |
| CHIT1 | chitinase 1 (chitotriosidase) |
| CHUK | conserved helix-loop-helix ubiquitous kinase |
| CKLF | chemokine-like factor |
| CLEC4A | C-type lectin domain family 4, member A |
| CLEC4C | C-type lectin domain family 4, member C |
| CLEC5A | C-type lectin domain family 5, member A |
| CLEC6A | C-type lectin domain family 6, member A |
| CLEC7A | C-type lectin domain family 7, member A |
| CLU | clusterin |
| CMA1 | chymase 1, mast cell |
| CMKLR1 | chemokine-like receptor 1 |
| COL3A1 | collagen, type III, alpha 1 |
| COLEC12 | collectin sub-family member 12 |
| CR1 |  |
| CR2 | complement component  (3d/Epstein Barr virus) receptor 2 |
| CREB1 | cAMP responsive element binding protein 1 |
| CREB5 | cAMP responsive element binding protein 5 |

| CREBBP | CREB binding protein |
| --- | --- |
| CRP | C-reactive protein, pentraxinrelated |
| CSF1 | colony stimulating factor 1 (macrophage) |
| CSF1R | colony stimulating factor 1 receptor |
| CSF2 | colony stimulating factor 2 (granulocyte-macrophage) |
| CSF2RB | colony stimulating factor 2 receptor, beta, low-affinity (granulocyte-macrophage) |
| CSF3 | colony stimulating factor 3 (granulocyte) |
| CSF3R | colony stimulating factor 3 receptor (granulocyte) |
| CT45A1 | cancer/testis antigen family 45, member A1 |
| CTAG1B | cancer/testis antigen 1B |
| CTAGE1 | cutaneous T-cell lymphomaassociated antigen 1 |
| CTCFL | CCCTC-binding factor (zinc finger protein)-like |
| CTLA4 | cytotoxic T-lymphocyte-associated protein 4 |
| CTSG | cathepsin G |
| CTSH | cathepsin H |
| CTSL | cathepsin L |
| CTSS | cathepsin S |
| CTSW | cathepsin W |
| CX3CL1 | chemokine (C-X3-C motif) ligand 1 |
| CX3CR1 | chemokine (C-X3-C motif) receptor 1 |
| CXCL1 | chemokine (C-X-C motif) ligand 1 (melanoma growth stimulating activity, alpha) |
| CXCL10 | chemokine (C-X-C motif) ligand 10 |
| CXCL11 | chemokine (C-X-C motif) ligand 11 |

| CXCL12 | chemokine (C-X-C motif) ligand 12 |
| --- | --- |
| CXCL13 | chemokine (C-X-C motif) ligand 13 |
| CXCL14 | chemokine (C-X-C motif) ligand 14 |
| CXCL16 | chemokine (C-X-C motif) ligand 16 |
| CXCL2 | chemokine (C-X-C motif) ligand 2 |
| CXCL3 | chemokine (C-X-C motif) ligand 3 |
| CXCL5 | chemokine (C-X-C motif) ligand 5 |
| CXCL6 | chemokine (C-X-C motif) ligand 6 (granulocyte chemotactic protein 2) |
| CXCL9 | chemokine (C-X-C motif) ligand 9 |
| CXCR1 | chemokine (C-X-C motif) receptor 1 |
| CXCR2 | chemokine (C-X-C motif) receptor 2 |
| CXCR3 | chemokine (C-X-C motif) receptor 3 |
| CXCR4 | chemokine (C-X-C motif) receptor 4 |
| CXCR5 | chemokine (C-X-C motif) receptor 5 |
| CXCR6 | chemokine (C-X-C motif) receptor 6 |
| CYBB | cytochrome b-245, beta polypeptide |
| CYFIP2 | cytoplasmic FMR1 interacting protein 2 |
| CYLD | cylindromatosis (turban tumor syndrome) |
| DDX43 | DEAD (Asp-Glu-Ala-Asp) box polypeptide 43 |
| DDX58 | DEAD (Asp-Glu-Ala-Asp) box polypeptide 58 |
| DEFB1 | defensin, beta 1 |
| DMBT1 | deleted in malignant brain tumors 1 |
| DOCK9 | dedicator of cytokinesis 9 |
| DPP4 | dipeptidyl-peptidase 4 |
| DUSP4 | dual specificity phosphatase 4 |

| DUSP6 | dual specificity phosphatase 6 |
| --- | --- |
| EBI3 | Epstein-Barr virus induced 3 |
| ECSIT | ECSIT homolog (Drosophila) |
| EGR1 | early growth response 1 |
| EGR2 | early growth response 2 |
| ELANE | elastase, neutrophil expressed |
| ELK1 | ELK1, member of ETS oncogene family |
| ENG | endoglin |
| ENTPD1 | ectonucleoside triphosphate diphosphohydrolase 1 |
| EOMES | eomesodermin |
| EP300 | E1A binding protein p300 |
| EPCAM | epithelial cell adhesion molecule |
| ETS1 | v-ets erythroblastosis virus E26 oncogene homolog 1 (avian) |
| EWSR1 | Ewing sarcoma breakpoint region 1 |
| F12 | coagulation factor XII (Hageman factor) |
| F13A1 | coagulation factor XIII, A1 polypeptide |
| F2RL1 | coagulation factor II (thrombin) receptor-like 1 |
| FADD | Fas (TNFRSF6)-associated via death domain |
| FAS | Fas (TNF receptor superfamily, member 6) |
| FCER1A |  |
| FCER1G | Fc fragment of IgE, high affinity I, receptor for; gamma polypeptide |
| FCER2 | Fc fragment of IgE, low affinity II, receptor for (CD23) |
| FCGR1A | Fc fragment of IgG, high affinity Ia, receptor (CD64) |
| FCGR2A | Fc fragment of IgG, low affinity IIa, receptor (CD32) |

| FCGR2B | Fc fragment of IgG, low affinity IIb, receptor (CD32) |
| --- | --- |
| FCGR3A | Fc fragment of IgG, low affinity IIIa, receptor (CD16a) |
| FEZ1 | fasciculation and elongation protein zeta 1 (zygin I) |
| FLT3 | fms-related tyrosine kinase 3 |
| FLT3LG | fms-related tyrosine kinase 3 ligand |
| FN1 | fibronectin 1 |
| FOS | FBJ murine osteosarcoma viral oncogene homolog |
| FOXJ1 | forkhead box J1 |
| FOXP3 | forkhead box P3 |
| FPR2 | formyl peptide receptor 2 |
| FUT5 | fucosyltransferase 5 (alpha (1,3) fucosyltransferase) |
| FUT7 | fucosyltransferase 7 (alpha (1,3) fucosyltransferase) |
| FYN | FYN oncogene related to SRC, FGR, YES |
| GAGE1 | G antigen 1 |
| GATA3 | GATA binding protein 3 |
| GNLY | granulysin |
| GPI | glucose-6-phosphate isomerase |
| GTF3C1 | general transcription factor IIIC, polypeptide 1, alpha 220kDa |
| GZMA | granzyme A (granzyme 1, cytotoxic T-lymphocyte-associated serine esterase 3) |
| GZMB | granzyme B (granzyme 2, cytotoxic T-lymphocyte-associated serine esterase 1) |
| GZMH | granzyme H (cathepsin G-like 2, protein h-CCPX) |
| GZMK | granzyme K (granzyme 3; tryptase  II) |
| GZMM | granzyme M (lymphocyte met-ase  1) |
| HAMP | hepcidin antimicrobial peptide |

| HAVCR2 | hepatitis A virus cellular receptor 2 |
| --- | --- |
| HCK | hemopoietic cell kinase |
| HLA-A | major histocompatibility complex, class I, A |
| HLA-B | major histocompatibility complex, class I, B |
| HLA-C | major histocompatibility complex, class I, C |
| HLA-DMA | major histocompatibility complex, class II, DM alpha |
| HLA-DMB | major histocompatibility complex, class II, DM beta |
| HLA-DOB | major histocompatibility complex, class II, DO beta |
| HLA-DPA1 | major histocompatibility complex, class II, DP alpha 1 |
| HLA-DPB1 | major histocompatibility complex, class II, DP beta 1 |
| HLA-DQA1 | major histocompatibility complex, class II, DQ alpha 1 |
| HLA-DQB1 | major histocompatibility complex, class II, DQ beta 1 |
| HLA-DRA | major histocompatibility complex, class II, DR alpha |
| HLA-DRB3 | major histocompatibility complex, class II, DR beta 3 |
| HLA-DRB4 | major histocompatibility complex, class II, DR beta 4 |
| HLA-E | major histocompatibility complex, class I, E |
| HLA-G | major histocompatibility complex, class I, G |
| HMGB1 | high mobility group box 1 |
| HRAS | v-Ha-ras Harvey rat sarcoma viral oncogene homolog |
| HSD11B1 | hydroxysteroid (11-beta) dehydrogenase 1 |
| ICAM1 | intercellular adhesion molecule 1 |
| ICAM2 | intercellular adhesion molecule 2 |
| ICAM3 | intercellular adhesion molecule 3 |
| ICAM4 | intercellular adhesion molecule 4 (Landsteiner-Wiener blood group) |

| ICOS | inducible T-cell co-stimulator |
| --- | --- |
| ICOSLG | inducible T-cell co-stimulator ligand |
| IDO1 | indoleamine 2,3-dioxygenase 1 |
| IFI16 | interferon, gamma-inducible protein 16 |
| IFI27 | interferon, alpha-inducible protein 27 |
| IFI35 | interferon-induced protein 35 |
| IFIH1 | interferon induced with helicase C domain 1 |
| IFIT1 | interferon-induced protein with tetratricopeptide repeats 1 |
| IFIT2 | interferon-induced protein with tetratricopeptide repeats 2 |
| IFITM1 | interferon induced transmembrane protein 1 |
| IFITM2 | interferon induced transmembrane protein 2 |
| IFNA1 | interferon, alpha 1 |
| IFNA17 | interferon, alpha 17 |
| IFNA2 | interferon, alpha 2 |
| IFNA7 | interferon, alpha 7 |
| IFNA8 | interferon, alpha 8 |
| IFNAR1 | interferon (alpha, beta and omega) receptor 1 |
| IFNAR2 | interferon (alpha, beta and omega) receptor 2 |
| IFNB1 | interferon, beta 1, fibroblast |
| IFNG | interferon, gamma |
| IFNGR1 | interferon gamma receptor 1 |
| IFNL1 | interferon lambda 1 |
| IFNL2 | interferon lambda 2 |
| IGF1R | insulin-like growth factor 1 receptor |
| IGF2R | insulin-like growth factor 2 receptor |

| IGLL1 | immunoglobulin lambda-like polypeptide 1 |
| --- | --- |
| IKBKB | inhibitor of kappa light polypeptide gene enhancer in B-cells, kinase beta |
| IKBKE | inhibitor of kappa light polypeptide gene enhancer in B-cells, kinase epsilon |
| IKBKG | inhibitor of kappa light polypeptide gene enhancer in B-cells, kinase gamma |
| IL17RB | interleukin 17 receptor B |

| IL18 | interleukin 18 (interferon-gammainducing factor) |
| --- | --- |
| IL18R1 | interleukin 18 receptor 1 |
| IL18RAP | interleukin 18 receptor accessory protein |
| IL19 | interleukin 19 |
| IL1A | interleukin 1, alpha |
| IL1B | interleukin 1, beta |
| IL1R1 | interleukin 1 receptor, type I |
| IL1R2 | interleukin 1 receptor, type II |
| IL1RAP | interleukin 1 receptor accessory protein |
| IL1RAPL2 | interleukin 1 receptor accessory protein-like 2 |
| IL1RL1 | interleukin 1 receptor-like 1 |
| IL1RL2 | interleukin 1 receptor-like 2 |
| IL1RN | interleukin 1 receptor antagonist |
| IL2 | interleukin 2 |
| IL21 | interleukin 21 |
| IL21R | interleukin 21 receptor |
| IL22 | interleukin 22 |
| IL22RA1 | interleukin 22 receptor, alpha 1 |
| IL22RA2 | interleukin 22 receptor, alpha 2 |
| IL23A | interleukin 23, alpha subunit p19 |
| IL23R | interleukin 23 receptor |
| IL24 | interleukin 24 |
| IL25 | interleukin 25 |
| IL26 | interleukin 26 |
| IL27 | interleukin 27 |
| IL2RA | interleukin 2 receptor, alpha |

| IL2RB | interleukin 2 receptor, beta |
| --- | --- |
| IL2RG | interleukin 2 receptor, gamma |
| IL3 | interleukin 3 (colony-stimulating factor, multiple) |
| IL32 | interleukin 32 |
| IL34 | interleukin 34 |
| IL3RA | interleukin 3 receptor, alpha (low affinity) |
| IL4 | interleukin 4 |
| IL4R | interleukin 4 receptor |
| IL5 | interleukin 5 (colony-stimulating factor, eosinophil) |
| IL5RA | interleukin 5 receptor, alpha |
| IL6 | interleukin 6 (interferon, beta 2) |
| IL6R | interleukin 6 receptor |
| IL6ST | interleukin 6 signal transducer (gp130, oncostatin M receptor) |
| IL7 | interleukin 7 |
| IL7R | interleukin 7 receptor |
| IL8 | C-X-C motif chemokine ligand 8 |
| IL9 | interleukin 9 |
| ILF3 | interleukin enhancer binding factor 3, 90kDa |
| INPP5D | inositol polyphosphate-5phosphatase, 145kDa |
| IRAK1 | interleukin-1 receptor-associated kinase 1 |
| IRAK2 | interleukin-1 receptor-associated kinase 2 |
| IRAK4 | interleukin-1 receptor-associated kinase 4 |
| IRF1 | interferon regulatory factor 1 |
| IRF2 | interferon regulatory factor 2 |
| IRF3 | interferon regulatory factor 3 |

| IRF4 | interferon regulatory factor 4 |
| --- | --- |
| IRF5 | interferon regulatory factor 5 |
| IRF7 | interferon regulatory factor 7 |
| IRF8 | interferon regulatory factor 8 |
| IRGM | immunity-related GTPase family, M |
| ISG15 | ISG15 ubiquitin-like modifier |
| ISG20 | interferon stimulated exonuclease gene 20kDa |
| ITCH | itchy E3 ubiquitin protein ligase |
| ITGA1 | integrin, alpha 1 |
| ITGA2 | integrin, alpha 2 (CD49B, alpha 2 subunit of VLA-2 receptor) |
| ITGA2B | integrin, alpha 2b (platelet glycoprotein IIb of IIb/IIIa complex, antigen CD41) |
| ITGA4 | integrin, alpha 4 (antigen CD49D, alpha 4 subunit of VLA-4 receptor) |
| ITGA5 | integrin, alpha 5 (fibronectin receptor, alpha polypeptide) |
| ITGA6 | integrin, alpha 6 |
| ITGAE | integrin, alpha E (antigen CD103, human mucosal lymphocyte antigen 1; alpha polypeptide) |
| ITGAL | integrin, alpha L (antigen CD11A (p180), lymphocyte functionassociated antigen 1; alpha polypeptide) |
| ITGAM | integrin, alpha M (complement component 3 receptor 3 subunit) |
| ITGAX | integrin, alpha X (complement component 3 receptor 4 subunit) |
| ITGB1 | integrin, beta 1 (fibronectin  receptor, beta polypeptide, antigen CD29 includes MDF2, MSK12) |
| ITGB2 | integrin, beta 2 (complement component 3 receptor 3 and 4 subunit) |
| ITGB3 | integrin, beta 3 (platelet glycoprotein IIIa, antigen CD61) |

| ITGB4 | integrin, beta 4 |
| --- | --- |
| ITK | IL2-inducible T-cell kinase |
| JAK1 | Janus kinase 1 |
| JAK2 | Janus kinase 2 |
| JAK3 | Janus kinase 3 |
| JAM3 | junctional adhesion molecule 3 |
| KIR_Activating_Subgro up_1 | killer cell immunoglobulin-like receptor, three domains, short cytoplasmic tail, 1 |
| KIR_Activating_Subgro up_2 | killer cell immunoglobulin-like receptor, two domains, short cytoplasmic tail, 1 |
| KIR_Inhibiting_Subgro up_1 | killer cell immunoglobulin-like receptor, two domains, long cytoplasmic tail, 1 |
| KIR_Inhibiting_Subgro up_2 | killer cell immunoglobulin-like receptor, two domains, long cytoplasmic tail, 3 |
| KIR3DL1 | killer cell immunoglobulin-like receptor, three domains, long cytoplasmic tail, 1 |
| KIR3DL2 | killer cell immunoglobulin-like receptor, three domains, long cytoplasmic tail, 2 |
| KIR3DL3 | killer cell immunoglobulin-like receptor, three domains, long cytoplasmic tail, 3 |
| KIT | v-kit Hardy-Zuckerman 4 feline sarcoma viral oncogene homolog |
| KLRB1 | killer cell lectin-like receptor subfamily B, member 1 |
| KLRC1 | killer cell lectin-like receptor subfamily C, member 1 |
| KLRC2 | killer cell lectin-like receptor subfamily C, member 2 |
| KLRD1 | killer cell lectin-like receptor subfamily D, member 1 |
| KLRF1 | killer cell lectin-like receptor subfamily F, member 1 |
| KLRG1 | killer cell lectin-like receptor subfamily G, member 1 |

| KLRK1 | killer cell lectin-like receptor subfamily K, member 1 |
| --- | --- |
| LAG3 | lymphocyte-activation gene 3 |
| LAIR2 | leukocyte-associated immunoglobulin-like receptor 2 |
| LAMP1 | lysosomal-associated membrane protein 1 |
| LAMP2 | lysosomal-associated membrane protein 2 |
| LAMP3 | lysosomal-associated membrane protein 3 |
| LBP | lipopolysaccharide binding protein |
| LCK | lymphocyte-specific protein tyrosine kinase |
| LCN2 | lipocalin 2 |
| LCP1 | lymphocyte cytosolic protein 1 (Lplastin) |
| LGALS3 | lectin, galactoside-binding, soluble, 3 |
| LIF | leukemia inhibitory factor |
| LILRA1 | leukocyte immunoglobulin-like receptor, subfamily A (with TM domain), member 1 |
| LILRA4 |  |
| LILRA5 |  |
| LILRB1 | leukocyte immunoglobulin-like receptor, subfamily B (with TM and ITIM domains), member 1 |
| LILRB2 | leukocyte immunoglobulin-like receptor, subfamily B (with TM and ITIM domains), member 2 |
| LILRB3 | leukocyte immunoglobulin-like receptor, subfamily B (with TM and ITIM domains), member 3 |
| LRP1 | low density lipoprotein receptorrelated protein 1 |
| LRRN3 | leucine rich repeat neuronal 3 |
| LTA | lymphotoxin alpha (TNF superfamily, member 1) |
| LTB | lymphotoxin beta (TNF superfamily, member 3) |

| LTBR | lymphotoxin beta receptor (TNFR superfamily, member 3) |
| --- | --- |
| LTF | lactotransferrin |
| LTK | leukocyte receptor tyrosine kinase |
| LY86 | lymphocyte antigen 86 |
| LY9 | lymphocyte antigen 9 |
| LY96 | lymphocyte antigen 96 |
| LYN | v-yes-1 Yamaguchi sarcoma viral related oncogene homolog |
| MAF | v-maf musculoaponeurotic  fibrosarcoma oncogene homolog (avian) |
| MAGEA1 | melanoma antigen family A, 1 (directs expression of antigen MZ2E) |
| MAGEA12 | melanoma antigen family A, 12 |
| MAGEA3 | melanoma antigen family A, 3 |
| MAGEA4 | melanoma antigen family A, 4 |
| MAGEB2 | melanoma antigen family B, 2 |
| MAGEC1 | melanoma antigen family C, 1 |
| MAGEC2 | melanoma antigen family C, 2 |
| MAP2K1 | mitogen-activated protein kinase kinase 1 |
| MAP2K2 | mitogen-activated protein kinase kinase 2 |
| MAP2K4 | mitogen-activated protein kinase kinase 4 |
| MAP3K1 |  |
| MAP3K5 | mitogen-activated protein kinase kinase kinase 5 |
| MAP3K7 | mitogen-activated protein kinase kinase kinase 7 |
| MAP4K2 | mitogen-activated protein kinase kinase kinase kinase 2 |
| MAPK1 | mitogen-activated protein kinase 1 |
| MAPK11 | mitogen-activated protein kinase 11 |

| MAPK14 | mitogen-activated protein kinase 14 |
| --- | --- |
| MAPK3 | mitogen-activated protein kinase 3 |
| MAPK8 | mitogen-activated protein kinase 8 |
| MAPKAPK2 | mitogen-activated protein kinaseactivated protein kinase 2 |
| MARCO | macrophage receptor with collagenous structure |
| MASP1 | mannan-binding lectin serine peptidase 1 (C4/C2 activating component of Ra-reactive factor) |
| MASP2 | mannan-binding lectin serine peptidase 2 |
| MAVS | mitochondrial antiviral signaling protein |
| MBL2 | mannose-binding lectin (protein C) 2, soluble |
| MIF | macrophage migration inhibitory factor (glycosylation-inhibiting factor) |

| MSR1 | macrophage scavenger receptor 1 |
| --- | --- |
| MST1R | macrophage stimulating 1 receptor (c-met-related tyrosine kinase) |
| MUC1 | mucin 1, cell surface associated |
| MX1 | myxovirus (influenza virus) resistance 1, interferon-inducible protein p78 (mouse) |
| MYD88 | myeloid differentiation primary response gene (88) |
| NCAM1 | neural cell adhesion molecule 1 |
| NCF4 | neutrophil cytosolic factor 4, 40kDa |
| NFATC4 | nuclear factor of activated T-cells, cytoplasmic, calcineurin-dependent 4 |
| NFKB1 | nuclear factor of kappa light polypeptide gene enhancer in B-cells 1 |
| NFKB2 | nuclear factor of kappa light polypeptide gene enhancer in B-cells 2 (p49/p100) |
| NFKBIA | nuclear factor of kappa light polypeptide gene enhancer in B-cells inhibitor,  alpha |
| NLRC5 | NLR family, CARD domain containing 5 |
| NLRP3 | NLR family, pyrin domain containing 3 |
| NOD1 | nucleotide-binding oligomerization domain containing 1 |

| NT5E | 5'-nucleotidase, ecto (CD73) |
| --- | --- |
| NUP107 | nucleoporin 107kDa |
| OAS3 | 2'-5'-oligoadenylate synthetase 3,  100kDa |
| OSM | oncostatin M |
| PASD1 | PAS domain containing 1 |
| PAX5 | paired box 5 |
| PBK | PDZ binding kinase |
| PDCD1 | programmed cell death 1 |
| PDCD1LG2 | programmed cell death 1 ligand 2 |
| PDGFC | platelet derived growth factor C |
| PDGFRB | platelet-derived growth factor receptor, beta polypeptide |
| PECAM1 | platelet/endothelial cell adhesion molecule 1 |
| PIK3CD |  |
| PIK3CG |  |
| PIN1 | peptidylprolyl cis/trans isomerase, NIMA-interacting 1 |
| PLA2G1B | phospholipase A2, group IB (pancreas) |
| PLA2G6 | phospholipase A2, group VI (cytosolic, calcium-independent) |
| PLAU | plasminogen activator, urokinase |
| PLAUR | plasminogen activator, urokinase receptor |
| PMCH | pro-melanin-concentrating hormone |
| PNMA1 | paraneoplastic Ma antigen 1 |
| POU2AF1 | POU class 2 associating factor 1 |
| POU2F2 | POU class 2 homeobox 2 |
| PPARG | peroxisome proliferator-activated receptor gamma |
| PPBP | pro-platelet basic protein  (chemokine (C-X-C motif) ligand 7) |

| PRAME | preferentially expressed antigen in melanoma |
| --- | --- |
| PRF1 | perforin 1 (pore forming protein) |
| PRG2 | proteoglycan 2, bone marrow  (natural killer cell activator, eosinophil granule major basic protein) |
| PRKCD | protein kinase C, delta |
| PRKCE | protein kinase C, epsilon |
| PRM1 | protamine 1 |
| PSEN1 | presenilin 1 |
| PSEN2 | presenilin 2 (Alzheimer disease 4) |
| PSMB10 | proteasome (prosome, macropain) subunit, beta type, 10 |
| PSMB7 | proteasome (prosome, macropain) subunit, beta type, 7 |
| PSMB8 | proteasome (prosome, macropain)  subunit, beta type, 8 (large multifunctional peptidase 7) |
| PSMB9 | proteasome (prosome, macropain)  subunit, beta type, 9 (large multifunctional peptidase 2) |
| PSMD7 | proteasome (prosome, macropain) 26S subunit, non-ATPase, 7 |
| PTGDR2 | prostaglandin D2 receptor 2 |
| PTGS2 | prostaglandin-endoperoxide synthase 2 (prostaglandin G/H synthase and cyclooxygenase) |
| PTPRC | protein tyrosine phosphatase, receptor type, C |
| PVR | poliovirus receptor |
| PYCARD | PYD and CARD domain containing |
| RAG1 | recombination activating gene 1 |
| REL | v-rel reticuloendotheliosis viral oncogene homolog (avian) |
| RELA | v-rel reticuloendotheliosis viral oncogene homolog A (avian) |
| RELB | v-rel reticuloendotheliosis viral oncogene homolog B |

| REPS1 | RALBP1 associated Eps domain containing 1 |
| --- | --- |
| RIPK2 | receptor-interacting serinethreonine kinase 2 |
| ROPN1 | rhophilin associated tail protein 1 |
| RORA | RAR-related orphan receptor A |
| RORC | RAR-related orphan receptor C |
| RPS6 | ribosomal protein S6 |
| RRAD | Ras-related associated with diabetes |
| RUNX1 | runt-related transcription factor 1 |
| RUNX3 | runt-related transcription factor 3 |
| S100A12 | S100 calcium binding protein A12 |
| S100A7 | S100 calcium binding protein A7 |
| S100A8 | S100 calcium binding protein A8 |
| S100B | S100 calcium binding protein B |
| SAA1 | serum amyloid A1 |
| SBNO2 | strawberry notch homolog 2 (Drosophila) |
| SELE | selectin E |
| SELL | selectin L |
| SELPLG | selectin P ligand |
| SEMG1 | semenogelin I |
| SERPINB2 | serpin peptidase inhibitor, clade B (ovalbumin), member 2 |
| SERPING1 | serpin peptidase inhibitor, clade G (C1 inhibitor), member 1 |
| SH2B2 | SH2B adaptor protein 2 |
| SH2D1A | SH2 domain containing 1A |
| SH2D1B | SH2 domain containing 1B |
| SIGIRR |  |
| SIGLEC1 | sialic acid binding Ig-like lectin 1, sialoadhesin |

| SLAMF1 | signaling lymphocytic activation molecule family member 1 |
| --- | --- |
| SLAMF6 | SLAM family member 6 |
| SLAMF7 | SLAM family member 7 |
| SLC11A1 | solute carrier family 11 (protoncoupled divalent metal ion transporters), member 1 |
| SMAD2 | SMAD family member 2 |
| SMAD3 | SMAD family member 3 |
| SMPD3 | sphingomyelin phosphodiesterase 3, neutral membrane (neutral sphingomyelinase II) |
| SOCS1 | suppressor of cytokine signaling 1 |
| SPA17 | sperm autoantigenic protein 17 |
| SPACA3 | sperm acrosome associated 3 |
| SPANXB1 | SPANX family, member B1 |
| SPINK5 | serine peptidase inhibitor, Kazal type 5 |
| SPN | sialophorin |
| SPO11 | SPO11 meiotic protein covalently bound to DSB homolog (S.  cerevisiae) |
| SPP1 | secreted phosphoprotein 1 |
| SSX1 | synovial sarcoma, X breakpoint 1 |
| SSX4 | synovial sarcoma, X breakpoint 4 |
| ST6GAL1 | ST6 beta-galactosamide alpha-2,6sialyltranferase 1 |
| STAT1 | signal transducer and activator of transcription 1, 91kDa |
| STAT2 | signal transducer and activator of transcription 2, 113kDa |
| STAT3 | signal transducer and activator of transcription 3 (acute-phase response factor) |
| STAT4 | signal transducer and activator of transcription 4 |
| STAT5B | signal transducer and activator of transcription 5B |

| STAT6 | signal transducer and activator of transcription 6, interleukin-4 induced |
| --- | --- |
| SYCP1 | synaptonemal complex protein 1 |
| SYK | spleen tyrosine kinase |
| SYT17 | synaptotagmin XVII |
| TAB1 | TGF-beta activated kinase  1/MAP3K7 binding protein 1 |
| TAL1 | T-cell acute lymphocytic leukemia 1 |
| TANK | TRAF family member-associated NFKB activator |
| TAP1 | transporter 1, ATP-binding cassette, sub-family B (MDR/TAP) |
| TAP2 | transporter 2, ATP-binding cassette, sub-family B (MDR/TAP) |
| TAPBP | TAP binding protein (tapasin) |
| TARP | TCR gamma alternate reading frame protein |
| TBK1 | TANK-binding kinase 1 |
| TBX21 | T-box 21 |
| TCF7 | transcription factor 7 (T-cell specific, HMG-box) |
| TFE3 | transcription factor binding to IGHM enhancer 3 |
| TFEB | transcription factor EB |
| TFRC | transferrin receptor (p90, CD71) |
| TGFB1 | transforming growth factor, beta 1 |
| TGFB2 | transforming growth factor, beta 2 |
| THBD | thrombomodulin |
| THBS1 | thrombospondin 1 |
| THY1 | Thy-1 cell surface antigen |
| TICAM1 | toll-like receptor adaptor molecule 1 |
| TICAM2 | toll-like receptor adaptor molecule 2 |

| TIGIT | T cell immunoreceptor with Ig and ITIM domains |
| --- | --- |
| TIRAP | toll-interleukin 1 receptor (TIR) domain containing adaptor protein |
| TLR1 | toll-like receptor 1 |

| TNFRSF17 | tumor necrosis factor receptor superfamily, member 17 |
| --- | --- |
| TNFRSF18 | tumor necrosis factor receptor superfamily, member 18 |
| TNFRSF1A | tumor necrosis factor receptor superfamily, member 1A |
| TNFRSF1B | tumor necrosis factor receptor superfamily, member 1B |
| TNFRSF4 | tumor necrosis factor receptor superfamily, member 4 |
| TNFRSF8 | tumor necrosis factor receptor superfamily, member 8 |
| TNFRSF9 | tumor necrosis factor receptor superfamily, member 9 |
| TNFSF10 | tumor necrosis factor (ligand) superfamily, member 10 |
| TNFSF11 | tumor necrosis factor (ligand) superfamily, member 11 |
| TNFSF12 | tumor necrosis factor (ligand) superfamily, member 12 |
| TNFSF13 | tumor necrosis factor (ligand) superfamily, member 13 |
| TNFSF13B | tumor necrosis factor (ligand) superfamily, member 13b |
| TNFSF14 | tumor necrosis factor (ligand) superfamily, member 14 |
| TNFSF15 | tumor necrosis factor (ligand) superfamily, member 15 |
| TNFSF18 | tumor necrosis factor (ligand) superfamily, member 18 |
| TNFSF4 | tumor necrosis factor (ligand) superfamily, member 4 |
| TNFSF8 | tumor necrosis factor (ligand) superfamily, member 8 |
| TOLLIP | toll interacting protein |
| TP53 | tumor protein p53 |
| TPSAB1 | tryptase alpha/beta 1 |
| TPTE | transmembrane phosphatase with tensin homology |
| TRAF2 | TNF receptor-associated factor 2 |
| TRAF3 | TNF receptor-associated factor 3 |
| TRAF6 | TNF receptor-associated factor 6, E3 ubiquitin protein ligase |

| TREM1 | triggering receptor expressed on myeloid cells 1 |
| --- | --- |
| TREM2 | triggering receptor expressed on myeloid cells 2 |
| TTK | TTK protein kinase |
| TXK | TXK tyrosine kinase |
| TXNIP | thioredoxin interacting protein |
| TYK2 | tyrosine kinase 2 |
| UBC | ubiquitin C |
| ULBP2 | UL16 binding protein 2 |
| USP9Y | ubiquitin specific peptidase 9, Ylinked |
| VCAM1 | vascular cell adhesion molecule 1 |
| VEGFA | vascular endothelial growth factor A |
| VEGFC | vascular endothelial growth factor C |
| XCL2 | chemokine (C motif) ligand 2 |
| XCR1 | chemokine (C motif) receptor 1 |
| YTHDF2 | YTH domain family, member 2 |
| ZAP70 | zeta-chain (TCR) associated protein kinase 70kDa |
| ZNF205 | zinc finger protein 205 |
| ABCF1 |  |
| AGK |  |
| ALAS1 |  |
| AMMECR1L |  |
| CC2D1B |  |
| CNOT10 |  |
| CNOT4 |  |
| COG7 |  |

| DDX50 | DEAD (Asp-Glu-Ala-Asp) box polypeptide 50 |
| --- | --- |
| DHX16 | DEAH (Asp-Glu-Ala-His) box polypeptide 16 |
| DNAJC14 | DnaJ (Hsp40) homolog, subfamily C, member 14 |
| EDC3 | enhancer of mRNA decapping 3 homolog (S. cerevisiae) |
| EIF2B4 | eukaryotic translation initiation factor 2B, subunit 4 delta, 67kDa |
| ERCC3 | excision repair cross-complementing rodent repair deficiency, complementation group 3 |
| FCF1 | FCF1 small subunit (SSU) processome component homolog (S. cerevisiae) |
| G6PD | glucose-6-phosphate dehydrogenase |
| GPATCH3 | G patch domain containing 3 |
| GUSB | glucuronidase, beta |
| HDAC3 | histone deacetylase 3 |
| HPRT1 | hypoxanthine phosphoribosyltransferase 1 |
| MRPS5 | mitochondrial ribosomal protein S5 |
| MTMR14 | myotubularin related protein 14 |
| NOL7 | nucleolar protein 7, 27kDa |
| NUBP1 | nucleotide binding protein 1 |
| POLR2A | polymerase (RNA) II (DNA directed) polypeptide A, 220kDa |
| PPIA | peptidylprolyl isomerase A (cyclophilin A) |
| PRPF38A | PRP38 pre-mRNA processing factor 38 (yeast) domain containing A |
| SAP130 | Sin3A-associated protein, 130kDa |
| SDHA | succinate dehydrogenase complex, subunit A, flavoprotein (Fp) |
| SF3A3 | splicing factor 3a, subunit 3, 60kDa |
| TBP | TATA box binding protein |
| TLK2 | tousled-like kinase 2 |
| TMUB2 | transmembrane and ubiquitin-like domain containing 2 |
| TRIM39 | tripartite motif containing 39 |
| TUBB | tubulin, beta class I |
| USP39 | ubiquitin specific peptidase 39 |
| ZC3H14 | zinc finger CCCH-type containing 14 |
| ZKSCAN5 | zinc finger with KRAB and SCAN domains 5 |
| ZNF143 | zinc finger protein 143 |
| ZNF346 | zinc finger protein 346 |

**Supplementary Table 2.** HHV8 genes included in the Nanostring nCounter analysis.

| **Name** | **Gene ID** |
| --- | --- |
| ORF73 | Gene ID: 4961527 |
| ORF50 | Gene ID: 4961526 |
| ORF71 | Gene ID: 4961494 |
| ORF72 | Gene ID: 4961471 |
| ORF74 | Gene ID: 4961460 |
| ORF16 | Gene ID: 4961447 |
| K2 | Gene ID: 4961449 |
| v-IRF1 | Gene ID: 4961464 |
| K8.1 | Gene ID: 4961469 |
| ORF75 | Gene ID: 4961476 |

**Supplementary Table 3.** List of Antibodies and BOND working conditions used for IHC and mIF panel 1.

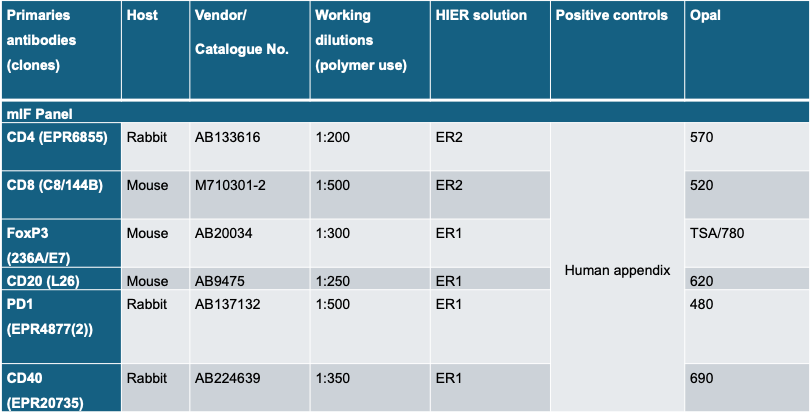

**Supplementary Table 4.** List of Antibodies and BOND working conditions used for IHC and mIF panel 2.

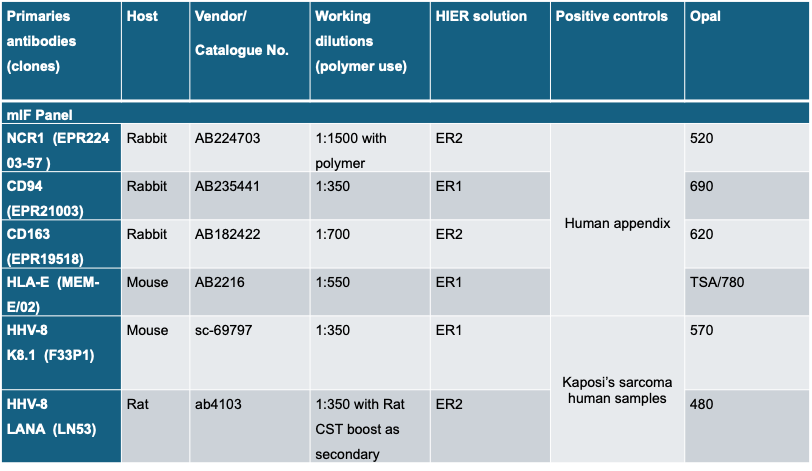

**Supplementary Table 5**. Top 30 genes differentially expressed in the 2 cohorts. Positive LogFC refers to genes that are upregulated in the non-HIV cohort, negative LogFC indicated genes that are upregulated in the HIV cohort.

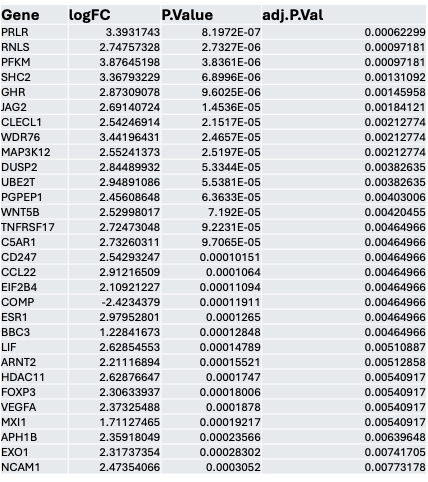

**Supplementary Table 6.** Host genes associated with the expression of HHV8 genes.

| **Gene** | **Spearman_rho** | **p_value** | **FDR** |
| --- | --- | --- | --- |
| NOS2 | 0.911755064 | 2.317E-10 | 1.74E-07 |
| RAD51 | 0.856923077 | 2.1874E-06 | 0.0002212 |
| JAK1 | 0.846153846 | 2.1215E-06 | 0.0002212 |
| CDH5 | 0.84 | 2.0926E-06 | 0.0002212 |
| BCAT1 | 0.826923077 | 2.1285E-06 | 0.0002212 |
| CD80 | 0.825384615 | 2.1479E-06 | 0.0002212 |
| TNFRSF10C | 0.82 | 2.2548E-06 | 0.0002212 |
| DNMT1 | 0.816153846 | 2.3766E-06 | 0.0002212 |
| BCL6B | 0.806153846 | 2.9421E-06 | 0.0002212 |
| H2AFX | 0.805384615 | 3.0043E-06 | 0.0002212 |
| BIRC5 | 0.802307692 | 3.2859E-06 | 0.0002212 |
| MYCT1 | 0.8 | 3.5345E-06 | 0.0002212 |
| BATF3 | 0.786153846 | 5.9061E-06 | 0.0003142 |
| CD40 | 0.783076923 | 6.6941E-06 | 0.0003142 |
| LILRA3 | 0.781538462 | 7.1321E-06 | 0.00031507 |
| DLL4 | 0.774615385 | 9.5181E-06 | 0.00038853 |
| TYMS | 0.773846154 | 9.8296E-06 | 0.00038853 |
| DEFB134 | 0.771485054 | 6.3281E-06 | 0.0003142 |
| IL21R | 0.766153846 | 1.3547E-05 | 0.00050871 |
| CTLA4 | 0.756153846 | 2.037E-05 | 0.00072847 |
| MMP9 | 0.752307692 | 2.3734E-05 | 0.00079883 |
| MELK | 0.746923077 | 2.927E-05 | 0.00090563 |
| FZD8 | 0.746153846 | 3.0147E-05 | 0.00090563 |
| MLH1 | 0.743076923 | 3.3891E-05 | 0.00097893 |
| CDC25C | 0.739030242 | 2.4465E-05 | 0.00079883 |
| LILRA1 | 0.738461538 | 4.0266E-05 | 0.00111999 |
| TNFRSF10B | 0.735384615 | 4.5073E-05 | 0.00116027 |
| TNFRSF10D | 0.735384615 | 4.5073E-05 | 0.00116027 |
| IL1B | 0.734615385 | 4.6349E-05 | 0.00116027 |
| PRF1 | 0.730769231 | 5.321E-05 | 0.00128906 |
| FANCA | 0.724615385 | 6.6008E-05 | 0.00154912 |
| IL10 | 0.717692308 | 8.3481E-05 | 0.00189983 |
| CCNB1 | 0.715384615 | 9.0124E-05 | 0.00199067 |
| HELLS | 0.707692308 | 0.00011564 | 0.0023344 |
| NFKBIE | 0.706923077 | 0.0001185 | 0.0023344 |
| PVR | 0.706153846 | 0.00012142 | 0.0023344 |
| UBE2C | 0.706153846 | 0.00012142 | 0.0023344 |
| CD300A | 0.704615385 | 0.00012744 | 0.0023344 |
| PLA1A | 0.704615385 | 0.00012744 | 0.0023344 |
| CENPF | 0.7 | 0.00014708 | 0.00262999 |
| STC1 | 0.699749965 | 9.8916E-05 | 0.00212245 |
| EIF5AL1 | 0.697692308 | 0.00015784 | 0.00263775 |
| ROBO4 | 0.696923077 | 0.00016157 | 0.00263775 |
| TNFRSF4 | 0.696923077 | 0.00016157 | 0.00263775 |
| BRCA2 | 0.695384615 | 0.00016925 | 0.00270445 |
| SELE | 0.694615385 | 0.00017321 | 0.00271005 |
| CDC20 | 0.693076923 | 0.00018137 | 0.00277976 |
| SNAI1 | 0.687692308 | 0.00021255 | 0.00312997 |
| POLD1 | 0.686153846 | 0.00022226 | 0.00321001 |
| FGF18 | 0.685516458 | 0.00015569 | 0.00263775 |
| BID | 0.683076923 | 0.00024283 | 0.0034408 |
| GZMB | 0.678976739 | 0.0001902 | 0.00285675 |
| FUT4 | 0.676153846 | 0.00029511 | 0.00402962 |
| PDGFA | 0.676153846 | 0.00029511 | 0.00402962 |
| CEP55 | 0.673846154 | 0.00031455 | 0.00412531 |
| VHL | 0.673846154 | 0.00031455 | 0.00412531 |
| EXO1 | 0.671538462 | 0.00033508 | 0.00412531 |
| MMP1 | 0.671538462 | 0.00033508 | 0.00412531 |
| RELB | 0.670769231 | 0.00034217 | 0.00414468 |
| IL24 | 0.667692308 | 0.00037184 | 0.00443251 |
| CXCL6 | 0.660511649 | 0.00032621 | 0.00412531 |
| ICAM2 | 0.658461538 | 0.0004744 | 0.00539807 |
| MAGEA3.A6 | 0.654831136 | 0.00038233 | 0.00448643 |
| LOXL2 | 0.653076923 | 0.00054473 | 0.00602746 |
| REN | 0.650097546 | 0.00043534 | 0.00502987 |
| SPRY4 | 0.643076923 | 0.00069919 | 0.00729294 |
| KIR2DL3 | 0.64166187 | 0.00054576 | 0.00602746 |
| FCAR | 0.641538462 | 0.000726 | 0.0074688 |
| PRKCA | 0.640892491 | 0.00055695 | 0.00606184 |
| IHH | 0.640769231 | 0.00073972 | 0.0075072 |
| GLS | 0.638584355 | 0.0005917 | 0.00634811 |
| OLR1 | 0.637692308 | 0.00079689 | 0.00787451 |
| SIGLEC5 | 0.636923077 | 0.00081175 | 0.00791725 |
| BMP2 | 0.636153846 | 0.00082686 | 0.00796117 |
| VEGFC | 0.633846154 | 0.00087362 | 0.00830495 |
| SIGLEC8 | 0.628967121 | 0.00075754 | 0.00758552 |
| FCGR3A.B | 0.628461538 | 0.00099163 | 0.00930889 |
| CCL20 | 0.627692308 | 0.00100955 | 0.00936013 |
| NRAS | 0.625384615 | 0.00106499 | 0.00975372 |
| CEACAM3 | 0.62 | 0.00120457 | 0.01064271 |
| CSF2 | 0.62 | 0.00120457 | 0.01064271 |
| GPC4 | 0.62 | 0.00120457 | 0.01064271 |
| IL6 | 0.614615385 | 0.00135951 | 0.01173553 |
| CXCL13 | 0.613846154 | 0.00138298 | 0.01180245 |
| CASP9 | 0.613076923 | 0.00140679 | 0.01187078 |
| FLT1 | 0.611538462 | 0.00145547 | 0.01211833 |
| MRE11 | 0.61 | 0.00150558 | 0.01229013 |
| NLRC5 | 0.607692308 | 0.00158351 | 0.0127873 |
| NCAM1 | 0.606923077 | 0.00161025 | 0.01286483 |
| FZD9 | 0.605081011 | 0.0013529 | 0.01173553 |
| CCL7 | 0.604615385 | 0.00169277 | 0.01338177 |
| FAM124B | 0.603076923 | 0.00174977 | 0.01368828 |
| MAGEC1 | 0.601548885 | 0.0014684 | 0.01211833 |
| RIPK2 | 0.601538462 | 0.0018084 | 0.01371826 |
| XCL1.2 | 0.601538462 | 0.0018084 | 0.01371826 |
| FADD | 0.600769231 | 0.00183834 | 0.01375877 |
| BRCA1 | 0.6 | 0.0018687 | 0.01375877 |
| TNFSF9 | 0.6 | 0.0018687 | 0.01375877 |
| ANLN | 0.596153846 | 0.00202702 | 0.01449802 |
| CCNE1 | 0.594615385 | 0.00209348 | 0.01455743 |
| MAGEB2 | 0.594615385 | 0.00209348 | 0.01455743 |
| KIF2C | 0.593191009 | 0.0017759 | 0.01371826 |
| ADORA2A | 0.593076923 | 0.00216179 | 0.01483386 |
| CDH2 | 0.592307692 | 0.00219665 | 0.01483386 |
| NCR1 | 0.592307692 | 0.00219665 | 0.01483386 |
| CD38 | 0.591538462 | 0.00223199 | 0.01483386 |
| ITGB3 | 0.590769231 | 0.00226782 | 0.01493974 |
| CCNA1 | 0.59045437 | 0.00188788 | 0.01376506 |
| ICAM5 | 0.59 | 0.00230413 | 0.01504698 |
| CSF3 | 0.589981827 | 0.00190782 | 0.01377666 |
| DTX4 | 0.588461538 | 0.00237825 | 0.01539712 |
| COL11A2 | 0.586153846 | 0.00249324 | 0.01586798 |
| HEY1 | 0.586153846 | 0.00249324 | 0.01586798 |
| PTGS2 | 0.584615385 | 0.0025725 | 0.01623487 |
| ARG2 | 0.583846154 | 0.00261293 | 0.01635261 |
| KIR3DL1 | 0.581538462 | 0.00273749 | 0.01697457 |
| KDR | 0.580769231 | 0.00278012 | 0.01697457 |
| TMEM140 | 0.580769231 | 0.00278012 | 0.01697457 |
| PFKFB3 | 0.577692308 | 0.00295634 | 0.01762072 |
| CD274 | 0.575384615 | 0.00309466 | 0.01815698 |
| E2F3 | 0.575384615 | 0.00309466 | 0.01815698 |
| LAG3 | 0.573846154 | 0.00318991 | 0.01857072 |
| KIR3DL2 | 0.570390637 | 0.00290914 | 0.01747812 |
| CCL4 | 0.57 | 0.00343901 | 0.01981302 |
| COL11A1 | 0.566923077 | 0.00365002 | 0.02076156 |
| IFITM2 | 0.566153846 | 0.00370446 | 0.02076156 |
| IL11 | 0.566153846 | 0.00370446 | 0.02076156 |
| MAPK10 | 0.565384615 | 0.00375959 | 0.02091444 |
| APLNR | 0.563846154 | 0.00387194 | 0.021225 |
| CSF2RB | 0.563846154 | 0.00387194 | 0.021225 |
| APH1B | 0.563076923 | 0.00392917 | 0.02138268 |
| HNF1A | 0.56203117 | 0.00345607 | 0.01981302 |
| SELL | 0.560769231 | 0.00410521 | 0.02191086 |
| KAT2B | 0.559230769 | 0.00422626 | 0.02212118 |
| TLR1 | 0.559230769 | 0.00422626 | 0.02212118 |
| PROM1 | 0.558461538 | 0.00428791 | 0.0222084 |
| CCL2 | 0.557692308 | 0.00435032 | 0.02237732 |
| ICOS | 0.555384615 | 0.00454219 | 0.02320536 |
| CD79B | 0.555106762 | 0.0039729 | 0.02146511 |
| UBE2T | 0.553846154 | 0.00467406 | 0.0237177 |
| MAGEC2 | 0.553352492 | 0.00411376 | 0.02191086 |
| RRM2 | 0.552307692 | 0.00480916 | 0.02423945 |
| MAGEA12 | 0.55180357 | 0.00424161 | 0.02212118 |
| MFNG | 0.551538462 | 0.00487794 | 0.02442221 |
| CDKN2A | 0.546153846 | 0.00538315 | 0.02659699 |
| CXCL1 | 0.543846154 | 0.00561284 | 0.02702078 |
| IL2 | 0.543846154 | 0.00561284 | 0.02702078 |
| INHBA | 0.541538462 | 0.00585077 | 0.02784309 |
| IFNG | 0.540769231 | 0.00593195 | 0.02784309 |
| MSH2 | 0.540769231 | 0.00593195 | 0.02784309 |
| PDGFB | 0.540769231 | 0.00593195 | 0.02784309 |
| GPR160 | 0.54 | 0.00601408 | 0.02805326 |
| PF4 | 0.539885395 | 0.00534246 | 0.02657077 |
| SLAMF7 | 0.538461538 | 0.00618123 | 0.02865496 |
| BLM | 0.538180429 | 0.00551803 | 0.02702078 |
| IFNA1 | 0.537863269 | 0.00555122 | 0.02702078 |
| LILRB2 | 0.535384615 | 0.00652732 | 0.03007372 |
| IL1A | 0.534615385 | 0.00661635 | 0.03029805 |
| MSH6 | 0.533076923 | 0.0067975 | 0.03075253 |
| GPSM3 | 0.532307692 | 0.00688962 | 0.03098268 |
| CD7 | 0.531538462 | 0.0069828 | 0.0312148 |
| ATF3 | 0.528461538 | 0.00736623 | 0.03273394 |
| ADGRE1 | 0.527168609 | 0.00677288 | 0.03075253 |
| CCL8 | 0.524615385 | 0.00787031 | 0.03476823 |
| CD19 | 0.523846154 | 0.00797453 | 0.03481901 |
| TAF3 | 0.520769231 | 0.00840311 | 0.03647822 |
| MTOR | 0.52 | 0.00851322 | 0.03659344 |
| BBC3 | 0.519230769 | 0.00862455 | 0.03659344 |
| FCGR1A | 0.519230769 | 0.00862455 | 0.03659344 |
| NID2 | 0.519230769 | 0.00862455 | 0.03659344 |
| SPP1 | 0.518461538 | 0.0087371 | 0.03686272 |
| TCL1A | 0.518176582 | 0.00796726 | 0.03481901 |
| TLR8 | 0.517692308 | 0.00885088 | 0.03699668 |
| CDK6 | 0.516923077 | 0.00896591 | 0.03699668 |
| IL15 | 0.516923077 | 0.00896591 | 0.03699668 |
| KLRD1 | 0.516923077 | 0.00896591 | 0.03699668 |
| TDO2 | 0.515384615 | 0.00919972 | 0.03775404 |
| MAGEA1 | 0.514615385 | 0.00931853 | 0.03803379 |
| SERPINA1 | 0.513076923 | 0.00956002 | 0.03880852 |
| WNT3A | 0.512307692 | 0.00968272 | 0.03909529 |
| PPARG | 0.510769231 | 0.00993208 | 0.03967551 |
| TIE1 | 0.51 | 0.01005877 | 0.03975861 |
| CXCL5 | 0.505003999 | 0.01003055 | 0.03975861 |
| NEIL1 | 0.501538462 | 0.01154399 | 0.04515383 |
| TNFRSF8 | 0.50038983 | 0.01085073 | 0.04266439 |
| BRIP1 | 0.5 | 0.01183279 | 0.0458063 |
| C5AR1 | 0.5 | 0.01183279 | 0.0458063 |
| SOCS1 | 0.498461538 | 0.01212759 | 0.0465524 |
| BAD | 0.497692308 | 0.01227728 | 0.04680323 |
| CES3 | 0.495384615 | 0.01273562 | 0.04830529 |
| TNFSF4 | 0.493846154 | 0.01304903 | 0.04899911 |
| CTAG1B | 0.493642535 | 0.01214949 | 0.0465524 |
| IL2RA | 0.493076923 | 0.01320813 | 0.04934978 |
| PIAS4 | 0.492307692 | 0.01336884 | 0.04970298 |
| CST2 | 0.490478947 | 0.01280123 | 0.04831017 |
| EGFR | -0.794615385 | 4.2595E-06 | 0.00024607 |
| ERBB2 | -0.672307692 | 0.00032811 | 0.00412531 |
| TPSAB1.B2 | -0.644615385 | 0.00067324 | 0.00712113 |
| CXCL14 | -0.595384615 | 0.00206002 | 0.01455743 |
| BNIP3L | -0.591538462 | 0.00223199 | 0.01483386 |
| RICTOR | -0.579230769 | 0.00286708 | 0.01736432 |
| SLC1A5 | -0.510769231 | 0.00993208 | 0.03967551 |

**Supplementary Tables 7.** Host genes associated with HHV8 lytic genes.

| Gene | Spearman_rho_Lytic | FDR_Lytic |
| --- | --- | --- |
| IHH | 0.684615385 | 0.00364065 |
| FAM124B | 0.675384615 | 0.004043 |
| CSF3 | 0.663970404 | 0.0040373 |
| COL11A2 | 0.657692308 | 0.00567875 |
| TNFSF9 | 0.642307692 | 0.00743161 |
| MRE11 | 0.633846154 | 0.00841142 |
| KIR3DL2 | 0.633509221 | 0.00713761 |
| MAPK10 | 0.623076923 | 0.00958387 |
| PF4 | 0.620039687 | 0.0088809 |
| PTGS2 | 0.607692308 | 0.01175894 |
| CDH2 | 0.606153846 | 0.01193844 |
| CCNA1 | 0.602771541 | 0.01131206 |
| ARG2 | 0.602307692 | 0.0127232 |
| E2F3 | 0.6 | 0.01308731 |
| MAGEC2 | 0.59943293 | 0.01157716 |
| KIR3DL1 | 0.599230769 | 0.01308731 |
| MAGEA12 | 0.597884008 | 0.01175894 |
| IL11 | 0.595384615 | 0.01357083 |
| IFNA1 | 0.587041552 | 0.01357083 |
| MFNG | 0.583846154 | 0.01649003 |
| APH1B | 0.579230769 | 0.01764898 |
| BBC3 | 0.578461538 | 0.01777621 |
| CXCL13 | 0.577692308 | 0.01790493 |
| CDKN2A | 0.576153846 | 0.01831213 |
| TAF3 | 0.575384615 | 0.01844519 |
| TNFRSF8 | 0.57374945 | 0.01682693 |
| CD79B | 0.566262753 | 0.01874082 |
| CD19 | 0.563076923 | 0.02169713 |
| BAD | 0.558461538 | 0.02267759 |
| WNT3A | 0.557692308 | 0.02268812 |
| HNF1A | 0.555106762 | 0.02169806 |
| ATF3 | 0.554615385 | 0.02354016 |
| CES3 | 0.554615385 | 0.02354016 |
| TCL1A | 0.553183315 | 0.02214142 |
| MAGEA1 | 0.552307692 | 0.02407785 |
| SELL | 0.552307692 | 0.02407785 |
| IL1A | 0.549230769 | 0.02498082 |
| SLAMF7 | 0.549230769 | 0.02498082 |
| CD7 | 0.545384615 | 0.02611188 |
| OTOA | 0.545384615 | 0.02611188 |
| BNIP3 | 0.541538462 | 0.02729148 |
| CTAG1B | 0.539885395 | 0.02588507 |
| IFNG | 0.538461538 | 0.02865496 |
| SPIB | 0.536153846 | 0.02930857 |
| IL2 | 0.535384615 | 0.02953022 |
| NEIL1 | 0.534615385 | 0.02975378 |
| CXCL5 | 0.527328871 | 0.03018768 |
| FGF9 | 0.521170285 | 0.03302868 |
| LTB | 0.520769231 | 0.03606133 |
| BLK | 0.519230769 | 0.03638786 |
| TLR7 | 0.517692308 | 0.03692785 |
| ADGRE1 | 0.515993251 | 0.03574748 |
| TLR9 | 0.515384615 | 0.03754886 |
| WDR76 | 0.515384615 | 0.03754886 |
| IL17A | 0.514714378 | 0.03614924 |
| TBX21 | 0.514615385 | 0.0378282 |
| MAGEA4 | 0.513944999 | 0.03638786 |
| C5AR1 | 0.512307692 | 0.03909529 |
| IL4 | 0.510983607 | 0.03752862 |
| BLM | 0.510098105 | 0.03754886 |
| WNT5B | 0.509230769 | 0.04044165 |
| SLC1A5 | -0.508461538 | 0.04044165 |
| PIAS4 | 0.507692308 | 0.04044165 |
| TNFSF4 | 0.507692308 | 0.04044165 |
| IL2RA | 0.506153846 | 0.04125759 |
| CST2 | 0.505866522 | 0.03969095 |
| NLRP3 | 0.503846154 | 0.04239851 |
| RELN | 0.503846154 | 0.04239851 |
| CD3E | 0.5 | 0.04465539 |
| RAD51C | 0.499230769 | 0.04498277 |
| PNOC | 0.493456651 | 0.04553457 |
| FAM30A | 0.493076923 | 0.04838686 |
| TDO2 | 0.492307692 | 0.04873787 |
| SREBF1 | -0.491538462 | 0.04909139 |

**Supplementary Tables 8.** Host genes associated with HHV8 latent genes.

| Gene | Spearman_rho_Latent | FDR_Latent |
| --- | --- | --- |
| PDGFB | 0.661538462 | 0.00620375 |
| CXorf36 | 0.638461538 | 0.00876825 |
| CCL2 | 0.636923077 | 0.00883519 |
| RRM2 | 0.632307692 | 0.00932106 |
| MKI67 | 0.603076923 | 0.01642594 |
| TIE1 | 0.588461538 | 0.02076821 |
| SERPINA1 | 0.586153846 | 0.02152209 |
| ANGPT2 | 0.576153846 | 0.0235981 |
| GNLY | 0.574615385 | 0.02383462 |
| NID2 | 0.572307692 | 0.02457282 |
| TLR8 | 0.571538462 | 0.02457282 |
| SPP1 | 0.566923077 | 0.02538115 |
| PLOD2 | 0.559230769 | 0.0275993 |
| MRC1 | 0.537692308 | 0.03507769 |
| FCGR1A | 0.536923077 | 0.03507769 |
| CCL3.L1 | 0.523846154 | 0.04247425 |
| CCL8 | 0.520769231 | 0.04444178 |
| PECAM1 | 0.519230769 | 0.04466924 |
| GIMAP6 | 0.512307692 | 0.0487516 |
| CRABP2 | -0.511538462 | 0.0487516 |
